# RNA virus-mediated metabolic reprogramming in a bloom-forming, marine diatom

**DOI:** 10.64898/2026.06.16.732687

**Authors:** Chana F. Kranzler, Daniel P. Lowenstein, Linoy Bamshid, Hiba Waldman Ben-Asher, Helen Fredricks, Ehud Zelzion, Jolie M. Tosten, Benjamin A.S. Van Mooy, Kimberlee Thamatrakoln

## Abstract

Diatoms are a widespread group of phytoplankton that disproportionately influence the global carbon cycle through substantial contributions to primary production and carbon export in the ocean. Marine viruses are major drivers of host metabolic reprogramming and mortality and it is increasingly clear that RNA viruses, which primarily infect eukaryotes, are abundant and distributed throughout the global ocean. Using an integrated, multi-omics approach, we characterized the molecular and metabolic response of the model, bloom-forming, centric diatom *Chaetoceros tenuissimus* to RNA virus infection. Time-resolved transcriptomics revealed coordinated, differential regulation of more than a third of host genes prior to host lysis, eliciting a cascade of metabolic responses that included early shifts in sulfur and lipid metabolism, induction of nitrogen assimilation pathways, and late-stage activation of stress, signaling and death-related genes. Lipidomics revealed substantial cellular enrichment of phosphatidylethanolamine, ceramide and triacylglycerol during RNA virus infection, as well as an infection-specific shift in fatty acid composition. Collectively, these findings provide insight into the intracellular requirements for RNA virus replication and illustrate a fundamental shift in resource partitioning in infected diatoms, advancing our understanding of how RNA virus infection transforms diatom host metabolic function and downstream ecosystem dynamics.

## Introduction

Marine phytoplankton are estimated to contribute roughly half of the primary production on the planet (1), generating organic carbon that forms the base of the marine food web, sustains the microbial loop and drives the biological carbon pump. With an estimated ∼10^7^ per milliliter seawater, marine viruses are regarded as agents of ecological and biogeochemical transformation in the ocean (2–4). Ecologically, viruses influence phytoplankton community structure and facilitate the collapse of phytoplankton blooms (5–8). Biogeochemically, viruses alter the fate of organic carbon in the ocean by shifting the balance between trophic transfer, surface ocean remineralization and export pathways (reviewed in 9). Virus-mediated lysis of phytoplankton catalyzes regenerative processes, circumventing the transfer of particulate organic matter to higher trophic levels, through the ‘viral shunt’ (3). Virus infection has also been linked to enhanced export via the ‘viral shuttle’, through enhanced particle aggregation and sinking of infected cells (10, 11).

Viral infection in phytoplankton also facilitates substantial reprogramming of host metabolism (12–17) towards an infected cell state, or ‘virocell’ (18), to maximize virus production. Many marine viral genomes encode auxiliary metabolic genes (AMGs) that augment or manipulate host metabolism (16, 19), conferring a degree of ‘independence’ from the host’s metabolic network. This transformation has the potential to fundamentally alter cellular function within a community, ultimately reshaping the quality and composition of organic matter released following virus-mediated host lysis (20–23).

Current estimates of virus-mediated impacts on marine phytoplankton communities are largely based on viruses with double-stranded (ds) DNA genomes, such as cyanophage and giant viruses (e.g. *Phycodnaviridae*) (16). However, it is increasingly evident that RNA viruses, which dominate eukaryotic viromes (24), may be as abundant (25–27) and ubiquitous (27–29) as dsDNA viruses in global ocean. Despite our growing knowledge of the ocean virosphere, relatively little is known about the specific and functional impacts of RNA viruses on marine ecosystems.

Multiple isolations studies (30, 31) and field observations of natural assemblages (32–36) indicate that RNA viruses are important pathogens of marine diatoms, one of the most widespread and globally significant groups of eukaryotic phytoplankton (37, 38) that contribute ∼40% of marine primary production (39). With cell walls comprised of amorphous biogenic silica, diatoms disproportionally ballast substantial vertical flux of particulate organic matter to the deep ocean (40–43). Diatom-infecting RNA viruses are from the family *Marnaviridae*, a group of positive sense, single-stranded (ss) RNA viruses in the order *Picornavirales* (30, 31). In striking contrast to the large dsDNA genomes (∼30-500 kb) of other well-studied phytoplankton viruses, *Marnaviridae* genomes are only ∼9 kb, encoding for replication-related machinery and a structural polyprotein (30). While recent work identified AMGs in some marine RNA viruses (29), no AMGs have been identified in diatom-infecting *Marnaviridae* isolates (30) and the cellular response (17, 44–46) and broader biogeochemical impacts (23, 32, 33) of virus infection in diatoms are still being elucidated. Here, using the RNA virus, CtenRNAV type-I (47) and the cosmopolitan, bloom-forming, marine, centric diatom, *Chaetoceros tenuissimus*, we characterized the metabolic reprogramming that occurs during virocell formation using a combination of transcriptomic and lipidomic profiling.

## Results and Discussion

### CtenRNAV virus infection dynamics in *C. tenuissimus*

Cultures of the diatom, *C. tenuissimus* (NIES 3715, formerly strain 2-10), were infected by the ssRNA virus, CtenRNAV type-I (47) alongside uninfected control cultures (Ctrl). A decrease in host concentration (Fig. 1A) and photosynthetic efficiency (F_v_/F_m_; Fig. 1B) were observed in infected cultures 4 days post infection (dpi) with near complete lysis of the cultures 8 dpi. Intracellular virus, quantified using quantitative reverse transcriptase PCR (qRT-PCR) targeting the virus-encoded replicase polyprotein, increased by more than two orders of magnitude within the first day of infection and ∼5 orders of magnitude by 4 dpi, indicating active viral replication (Fig. 1C). Intracellular virus concentration (viruses mL^-1^) then decreased ∼100-fold between 4 and 8 dpi (Fig. 1C), coinciding with the onset of culture demise. Extracellular infectious virus production, measured using the most probable number (MPN) assay, was apparent by ∼2 dpi, increasing nearly two orders of magnitude compared to the initial virus titer. Extracellular virus concentration continued to increase until 5 dpi, reaching a final maximum concentration of ∼10^10^ viruses mL^-1^ (Fig. 1D). Biomass samples for RNA sequencing and particulate lipid analysis were collected 2 hours post infection (0.08 dpi) and then daily from 1-4 dpi, prior to host lysis.

**Figure 1.**
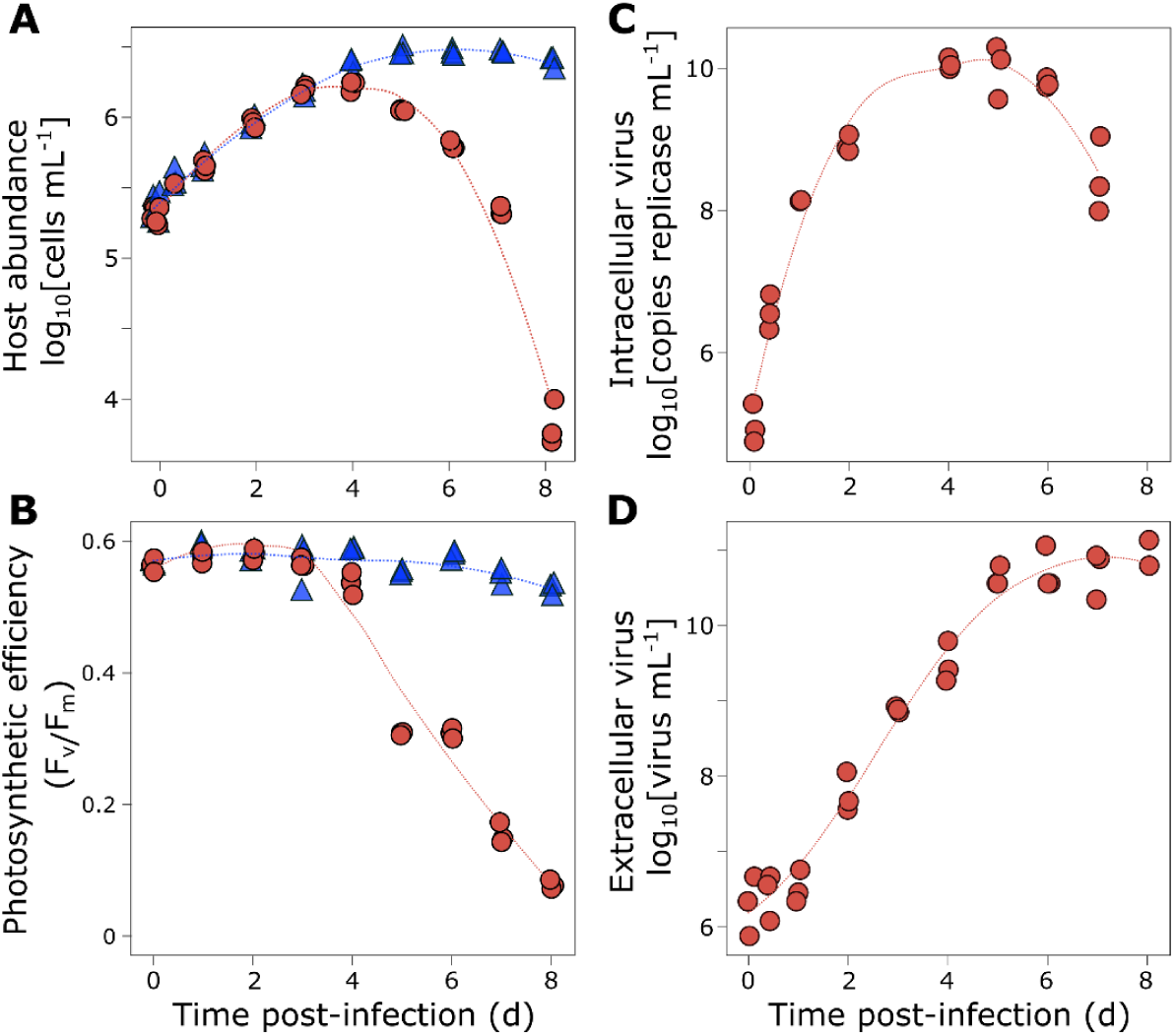
Host-virus dynamics during CtenRNAV virus infection. **A)** Host concentration (cells mL^-1^) and **B)** photosynthetic efficiency (F_v_/F_m_) in uninfected Ctrl (blue triangles) and CtenRNAV-infected *C. tenuissimus* cultures (red circles). **C)** qRT-PCR quantification of intracellular virus (copies replicase mL^-1^) and **D)** MPN quantification of extracellular infectious virus (virus mL^-1^) throughout the course of infection. Biological triplicates are plotted with LOESS regression. Data are shown for a single experiment and are representative of 2-3 infection experiments.

In infected treatments, RNA-seq reads mapping to the CtenRNAV genome increased steadily throughout the time course of infection (Fig. S1), consistent with the increase in intracellular virus measured by qRT-PCR (Fig. 1C). Read abundance for both the viral replicase polyprotein (D1P48_gp1) and the structural polyprotein (D1P48 gp2) increased by more than ∼2 orders of magnitude by 1 dpi, further increasing an additional ∼2 orders of magnitude by 4 dpi (Fig. S1). Mapped reads for the viral replicase polyprotein (D1P48_gp1) and structural polyprotein (D1P48_gp2) genes were also tightly correlated throughout the time course of infection (r^2^ = 0.999, *P* < 0.001; Fig. S1).

### Transcriptomic reprogramming in *C. tenuissimus* during CtenRNAV virus infection

After trimming raw RNA-seq data, 7.6-15.6 million reads (median 10.6 million) were obtained across the 30 samples (Data S1). Of these reads, ∼83% mapped to the *C. tenuissimus* reference genome (GenBank Accession No. GCA_021927905.1) and ∼69% of uniquely mapped reads were found within exons (Data S1). To expand the functional characterization of the 18,866 predicted proteins in the *C. tenuissimus* genome (48), we performed additional annotation (see Methods; Data S2). More than 50% (10,538) of the total predicted protein sequences were assigned a robust ortholog using a reciprocal blast in the UniProt database (49). For 5,601 of these predicted proteins (∼30% of the proteome, ∼53% of identified orthologs), 66% were assigned a Kegg ID and 53% were assigned at least one GO term. (Fig. S2; Data S2). Together, we identified 6,483 and 6,712 shared orthologs with *Phaeodactylum tricornutum* and *Thalassiosira pseudonana* diatom reference genomes, respectively, with 5,174 common genes across the three genomes (Fig. S2).

Infected cultures exhibited a significant shift (|log_2_FC| > 1; FDR < 0.05) in the expression of ∼35% of the 18,866 predicted genes in *C. tenuissimus* throughout the time course of infection (Data S3). Of those, 264 were differentially expressed 0.08 dpi compared to uninfected Ctrl cells (Fig. 2A; Fig. S3), with the overwhelming majority of differentially expressed genes (DEGs) downregulated (93%; 246). By 1 and 2 dpi, 1,643 and 1,179 DEGs, respectively, were detected in infected cells compared to Ctrl cells, with 51-62% upregulated. By 3 dpi, 71% of the 3,693 DEGs were upregulated during infection. The largest number of DEGs was detected 4 dpi, with 6,646 DEGs detected in infected cells; 58% of which were upregulated (Fig. 2A; Fig. S3).

**Figure 2.**
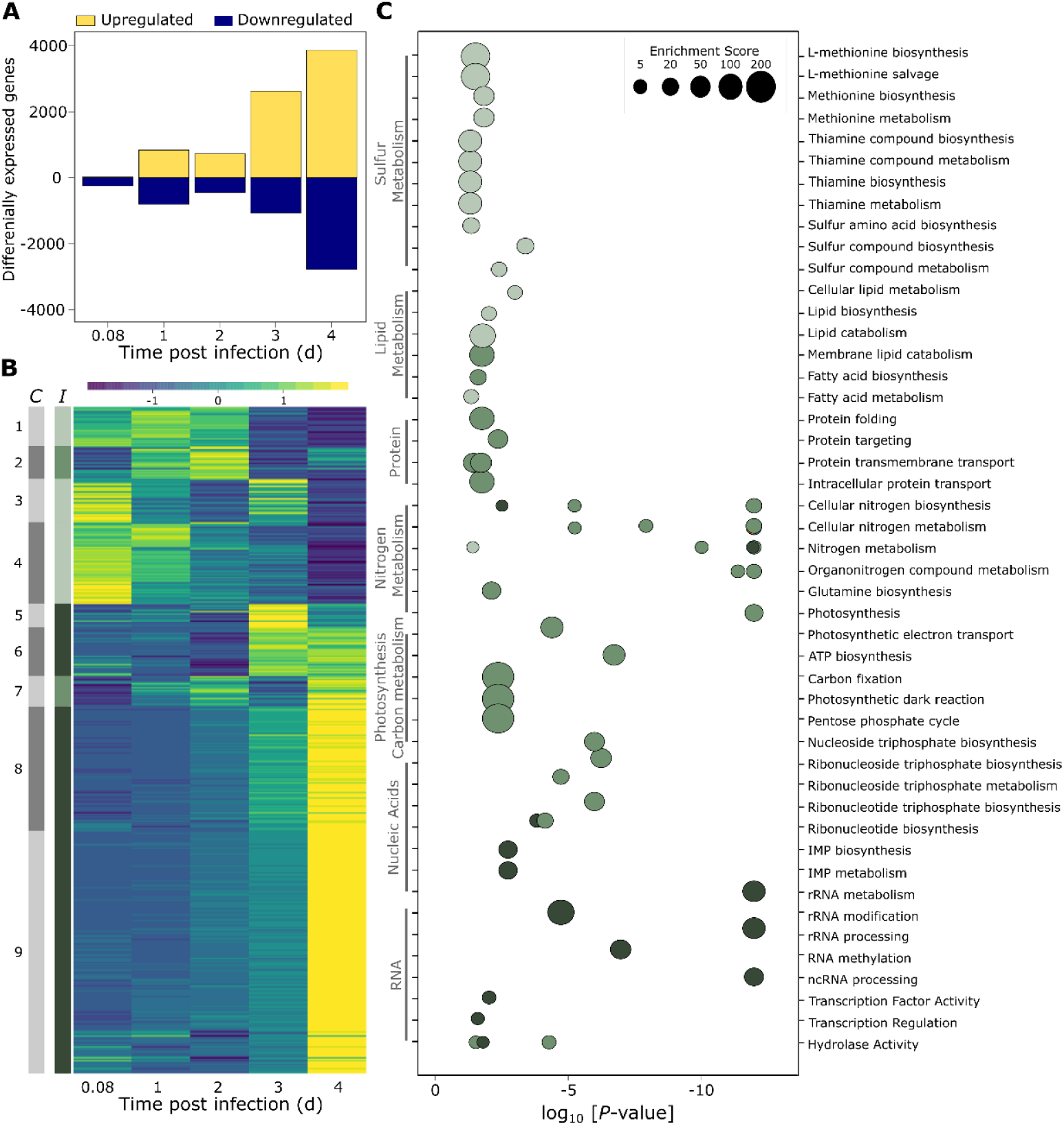
Differential expression, clustering and functional enrichment analysis of host genes expressed during CtenRNAV infection. **A)** Total number of upregulated (yellow) and downregulated (blue) differentially expressed genes in *C. tenuissimus* (|log_2_FC| > 1, FDR < 0.05) during the time course of CtenRNAV infection. **B**) Heatmap of scaled expression of differentially expressed genes, clustered by expression dynamics (*C*; K-means, n = 9) and stage of infection (*I*): ‘Early’ (light green), ‘Mid’ (green) and ‘Late’ (dark green). Heatmap color bar indicates scaled expression values. **C**) Gene Ontology (GO) terms that were significantly enriched within each cluster (*P* < 0.05, two-tailed, hyper geometric function). Symbol size denotes GO term enrichment score and colors denote temporal stages of infection as depicted in panel B. For visualization, genes with *P*-values < 10^-12^ are positioned at a value of 10^-12^. A complete list of all enriched GO terms can be found in Data S5.

Based on the scaled expression of DEGs in infected *C. tenuissimus*, nine distinct clusters of co-expressed genes were identified during the time course of infection, each exhibiting a distinct expression pattern (Fig. 2B; Fig. S4; Data S4). Transcripts in clusters 1, 3, and 4 were already highly expressed during the early stages of infection (0.08-1 dpi; Fig. 2B; Fig. S4). After this early peak, gene expression in cluster 4 steadily decreased, while those in cluster 1 remained elevated until 2 dpi and then declined. Expression of genes in cluster 3 decreased between 0.08 and 2 dpi but then increased again to peak expression 3 dpi before decreasing again. Expression of genes in cluster 2 and 7 increased from 0.08 dpi to a peak at 2 dpi, decreased from 2 to 3 dpi and then increased again by 4 dpi (Fig. 2B; Fig. S4). In clusters 5, 6, 8, and 9, genes were not strongly expressed until 3-4 dpi (Fig. 2B; Fig. S4). Based on these distinct expression patterns, clusters were categorized by the timing of the host response to the progression of CtenRNAV infection: ‘Early’ (clusters 1, 3 and 4), ‘Mid’ (clusters 2 and 7) and ‘Late’ (clusters 5, 6, 8 and 9; Fig. 2B; Fig. S4).

Functional enrichment analysis of each cluster based on Gene Ontology (GO) showed distinct metabolic shifts in *C. tenuissimus* corresponding to the stage of CtenRNAV infection (Fig. 2C, Data S5). ‘Early’ clusters were enriched in genes involved in lipid metabolism, fatty acid synthesis and lipid catabolism (Fig. 2C; Data S5), suggesting an immediate shift in cellular lipids in response to infection. GO terms related to methionine salvage, metabolism and biosynthesis and thiamine biosynthesis and metabolism pathways were also enriched during the early phase of infection (Fig. 2C), suggesting a shift in sulfur metabolism in infected cells. Interestingly, virus-induced modification of methionine cycling in plants can suppress host antiviral defense and promote infection (50), suggesting that a similar mechanism may operate during virus infection of diatoms.

During ‘Mid’ infection, clusters were also enriched in genes involved in membrane lipid catabolism and fatty acid biosynthesis (Fig. 2C; Data S5), suggesting substantial and prolonged modification of host lipid metabolism during CtenRNAV infection. Additional enriched categories during ‘Mid’ infection included protein homeostasis-related categories such as protein folding, targeting and transport, and genes involved in nitrogen metabolism, photosynthesis, carbon metabolism, and nucleotide biosynthesis (Fig. 2C). Genes expressed during ‘Late’ infection also exhibited enrichment in nitrogen metabolism-related genes and those related to RNA metabolism, including ribosomal RNA processing and modification, RNA methylation, non-coding RNA processing and transcriptional regulation (Fig. 2C; Data S5). Enrichment in genes with hydrolase activity was observed during both ‘Mid’ and ‘Late’ infection, including genes involved in cellular stress response and DNA repair (Fig. 2C).

#### Virus-induced shifts in cell signaling, stress and death

Throughout CtenRNAV infection, there was a marked shift in expression among genes involved in signaling, cellular stress, and death (Fig. 3). Several calmodulins (CaM), intracellular calcium sensors, were differentially expressed during CtenRNAV infection, including one CaM (GFH46458.1) that exhibited 6.5-fold higher expression in infected cells compared to Ctrl by 2 dpi, which increased to 21- and 67-fold by 3 and 4 dpi, respectively (Fig. 3A), suggesting a virus-induced shift in calcium homeostasis. As early as 1 dpi, a gene encoding for an additional EF-hand domain-containing protein (GFH53108.1) exhibited a 3-fold increase in expression, which was further upregulated 27-, 61- and 55-fold by 2, 3 and 4 dpi, respectively (Fig. 3A). These shifts were concomitant with ∼3-fold reduced expression of two calcium antiporters (GFH58649.1, GFH47508.1) by 3-4 dpi (Data S3), further suggesting a shift in internal calcium mobilization during CtenRNAV infection. In diatoms, calcium-mediated signaling has been implicated in stress surveillance and the activation of programmed cell death (PCD) pathways (51–56). Calcium homeostasis is also intricately involved in multiple aspects of the viral life cycle across diverse model systems (57). For example, in other members of *Picornavirales* such as *Coxsackievirus*, a virus-mediated reduction in calcium levels within subcellular compartments, such as the Golgi complex and ER, was shown to inhibit apoptosis, enabling virion assembly prior to host lysis (58).

**Figure 3.**
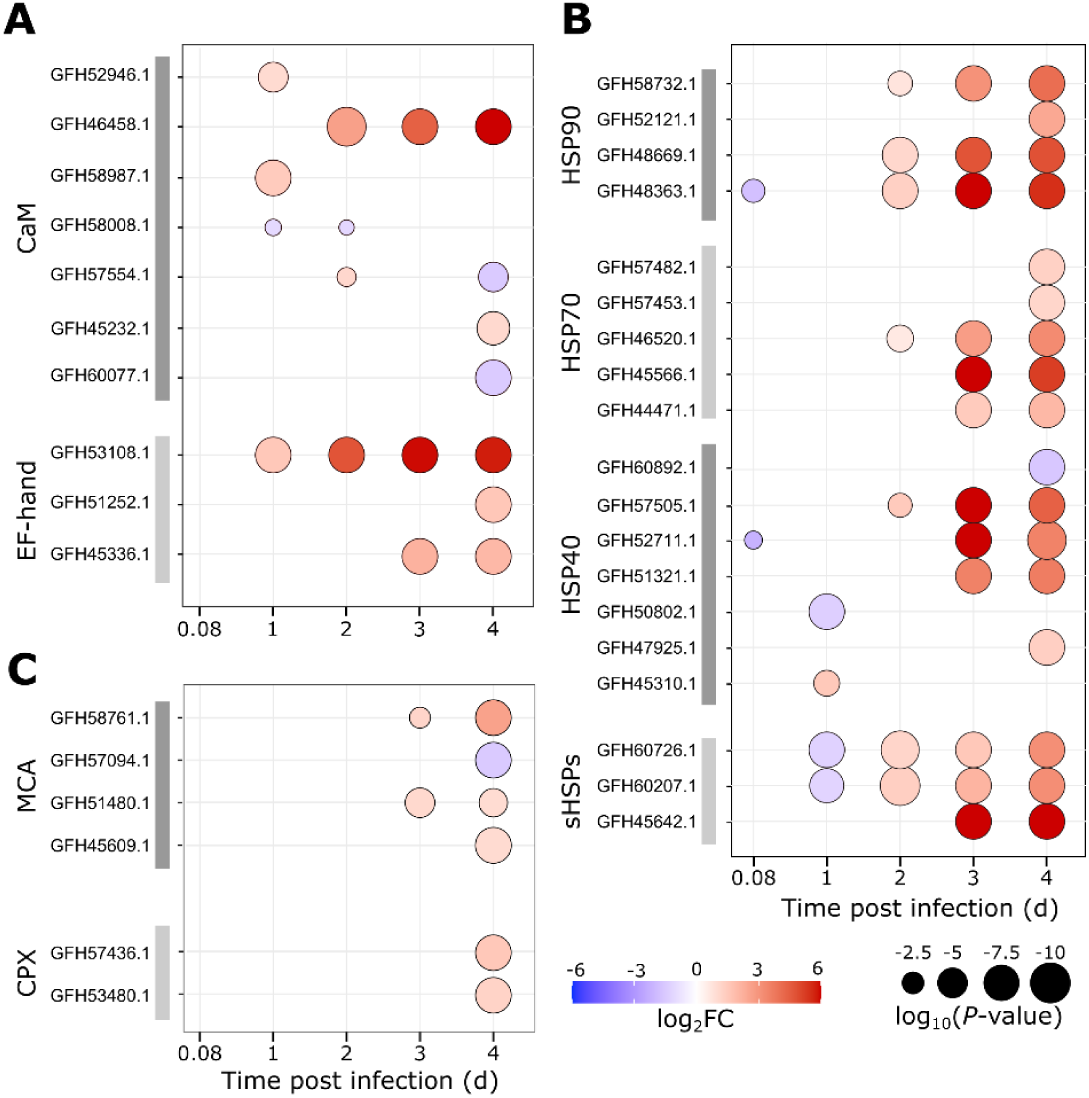
Modifications in stress, signaling and programmed cell death-related host gene expression during CtenRNAV infection. Differential expression analysis of genes related to **A**) calcium signaling, including calmodulin (CaM) and additional EF-hand domain-containing proteins, **B)** heat shock proteins (HSPs), and **C**) metacaspases (MCA) and cathepsin X (CPX) in infected *C. tenuissimus* compared to uninfected, Ctrl cultures. Symbol color depicts Fold Change (log_2_FC) and symbol size depicts adjusted *P*-values [log_10_(*P*-value)] for each pairwise comparison. Only significantly differentially expressed genes are shown (|log_2_FC| > 1; FDR < 0.05).

Several genes encoding heat shock proteins (HSPs) were also downregulated during early infection, with HSP90 (GFH48363.1) and HSP70 (GFH52711.1) exhibiting a 3.5-fold and 4.6-fold downregulation, respectively, 0.08 dpi in CtenRNAV-infected cultures compared to uninfected Ctrl cultures (Fig. 3B). Several additional HSPs were downregulated ∼2-3-fold 1 dpi (Fig. 3B). However, this expression pattern shifted substantially as the infection progressed, with significant upregulation of multiple HSPs by 2-4 dpi in CtenRNAV-infected cultures, including genes encoding HSP90, HSP70, HSP40 and small HSPs (sHSP; Fig. 3B). Notably, several HSPs were among the most highly upregulated DEGs during late CtenRNAV infection, exhibiting ∼135-210-fold increased expression (Fig. 3B, Data S3). As highly conserved, stress-inducible molecular chaperones, HSPs play an essential role in protein folding and stabilization (59, 60). In diatoms, HSPs are upregulated by a range of environmental stress conditions such high light (61), nutrient and trace metal deficiency (62, 63) and senescence (64). In addition, HSPs have been shown to interface with multiple stages of the viral life cycle in diverse model systems, including viral entry, replication and assembly (65).

The induction of autophagy genes (ATG) in *C. tenuissuimus* was recently reported during infection by CtenRNAV type-II (17), a taxonomically distinct virus (family *Marnaviridae*, genus *Sogarnavirus*; 65, 66). We also observed increased expression (2-11-fold) of multiple autophagy-related genes (ATGs) during CtenRNAV type-I infection, including ATG1 (GFH61212.1), ATG13 (GFH52049.1), ATG9 (GFH48833.1), ATG18 (GFH47286.1), ATG7 (GFH60098.1) and ATG12 (GFH54902.1) during late CtenRNAV infection (3-4 dpi; Data S3).

In other phytoplankton host-virus models systems, such as *Geophyrocapsa huxleyi (*formerly *Emiliania huxleyi)*, lytic virus infection triggers signatures of programmed cell death (PCD), including increased reactive oxygen species (ROS) and the induction of metacaspases (6, 68), a group of cysteine-dependent proteases associated with PCD in diverse groups of phytoplankton (56, 69–71). Consistent with these observations, several antioxidant-encoding genes were substantially upregulated by late CtenRNAV infection, including a peroxiredoxin (GFH55061.1; ∼100-fold) and a peroxidase (GFH61364.1; ∼30-fold; Data. S3), indicating a virus-mediated shift in ROS metabolism. This is consistent with prior work documenting increased intracellular ROS during advanced stages of virus infection in *C. tenuissimus* (33). We also found significant remodeling of genes associated with PCD during late-stage CtenRNAV infection (3-4 dpi), concurrent with the onset of culture demise (Fig. 3C). Out of six putative metacaspases (MCA) encoded in the *C. tenuissimus* genome, three were upregulated ∼2 - 6-fold at 3-4 dpi, with one MCA (GFH57094.1) downregulated ∼3-fold by 4 dpi (Fig. 3C). Two genes encoding for cathepsin X (CPX) were also ∼2-3-fold upregulated at 4 dpi (Fig. 3C). CPX, a cysteine protease, was recently shown to be a conserved marker for PCD onset in diatoms (72). Together, these expression patterns are consistent with virus-mediated onset of cellular stress and death in *C. tenuissimus,* indicating a shift in calcium signaling, ROS metabolism and the induction of a PCD-like pathway during CtenRNAV infection.

#### Virus-mediated modifications in cellular nitrogen metabolism

Throughout CtenRNAV infection in *C. tenuissimus*, there was substantial transcriptomic remodeling in nitrogen acquisition and assimilation genes prior to the onset of virus-mediated mortality and culture collapse (Fig. 4). Nitrate availability limits primary production in vast swaths of the ocean (73) and diatoms rely on a diverse suite of specialized nitrogen transporters, including those for nitrate (NO₃⁻), ammonium (NH₄⁺) and urea to assimilate nitrogen via a number of tightly regulated pathways (74). Three nitrate transporters (NRT) were upregulated in infected cells with two (NRT2; GFH58894.1 and GFH57077.1) upregulated ∼3-4-fold by 1 dpi, increasing 44-fold and 110-fold by 4 dpi, respectively and an NRT3 (GFH46206.1) increasing 7- and 15-fold at 3 and 4 dpi, respectively (Fig. 4). Nitrate reductase (NR; GFH58644.1) was among the few genes already upregulated by 0.08 dpi in infected cells, exhibiting ∼2-fold higher expression than uninfected Ctrl cells that progressively increased ∼6-, ∼21- and ∼77-fold at 1, 3 and 4 dpi, respectively (Fig. 4). In addition to a ∼4-fold increase in expression of a formate-nitrite transporter (FNT, GFH48708.1) at 1 dpi, we also observed upregulation in two nitrite reductases (NIR; GFH53591.1, NIRA; GFH52229.1) during the course CtenRNAV infection, with NIR expression increasing ∼4-30-fold at 2-4 dpi and NIRA exhibiting ∼3, 12- and 10-fold increased expression at 1, 3 and 4 dpi, respectively (Fig. 4).

**Figure 4.**
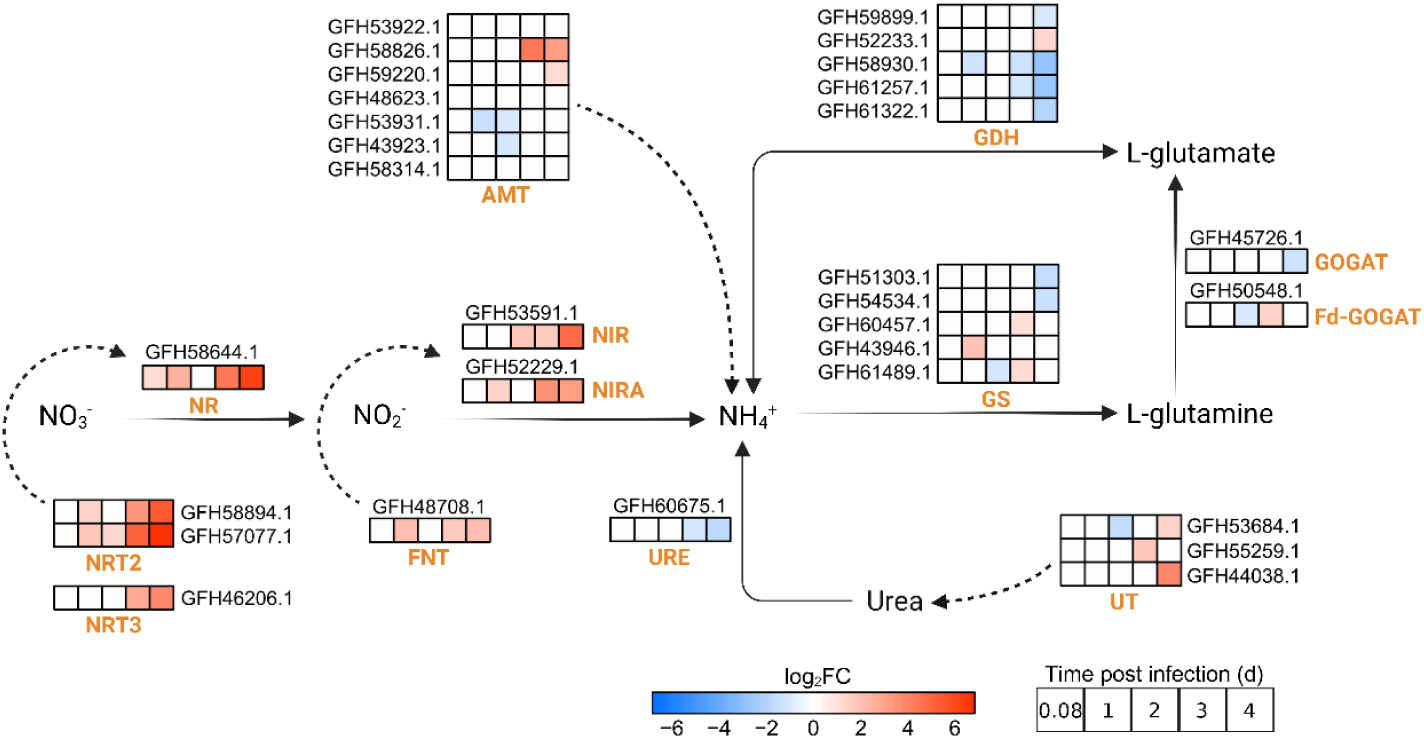
Modification in nitrogen assimilation pathways in infected *C. tenuissimus*. Schematic of genes involved in nitrogen assimilation throughout the time course of CtenRNAV virus infection. Heatmap color scale represents log_2_Fold Change (log_2_FC) values for DEGs in infected cultures compared to uninfected Ctrl cultures at each timepoint (individual boxes). White boxes represent timepoints in which genes were not significantly differentially expressed. Abbreviations are as follows: nitrate transporter (NRT), nitrate reductase (NR), formate-nitrite transporter (FNT), nitrite reductase (NIR), ferredoxin-nitrite reductase (NIRA), ammonium transporter (AMT), glutamate dehydrogenase (GDH), glutamine synthetase (GS), glutamate synthase (GOGAT), ferredoxin glutamate synthase (Fd-GOGAT), urease (URE), urea transporter (UT). Full description of gene annotations, fold change and adjusted *P*-values (FDR) are available in Data S2 and S3.

Among genes involved in the glutamine synthetase/glutamate synthase (GS/GOGAT) pathway, responsible for incorporating inorganic nitrogen into organic compounds, one of five putative GSs (GFH43946.1) was ∼4-fold upregulated at 1 dpi, suggesting enhanced ammonium assimilation during an early stage of CtenRNAV infection (Fig. 4). Two additional GSs (GFH60457.1 and GFH61489.1) were upregulated ∼2-fold at 3 dpi, while the other two (GFH51303.1 and GFH54534.1) were downregulated ∼3-fold by 4 dpi (Fig. 4). The majority of GOGAT genes and glutamate dehydrogenases (GDH) were also ∼2-7-fold downregulated at 3-4 dpi (Fig. 4), suggesting an additional shift in nitrogen assimilation during late-stage virus infection. Additionally, two out of seven ammonium transporters (AMT; GFH58826.1 and GFH59220.1) and all three urea transporters (UT; GFH53684.1, GFH55259.1 and GFH44038.1) were upregulated during late infection (3-4 dpi; Fig. 4). Nine out of 23 putative amino acid transporters also exhibited ∼2-55-fold increased expression at 1 to 4 dpi, with the most pronounced upregulation apparent within seven transporters at 3-4 dpi (Fig. S5).

The documented upregulation of nitrogen assimilation genes (NRT, NR, AMT) during CtenRNAV infection is consistent with the transcriptomic response to nitrogen limitation in diatoms (75–77). Despite the absence of virus-encoded AMGs in diatom-infecting RNA viruses that could enhance nutrient transport (78, 79), these findings suggest CtenRNAV infection may rewire host transcriptional networks to augment nitrogen uptake. This may be due to the stoichiometric mismatch between host biomass and virus requirements; i.e. the nitrogen-rich viral capsid and relatively high nucleic acid concentration require a lower C:N:P ratio than typical diatom biomass (80, 81), generating selective pressure on virocells to manipulate host nutrient acquisition pathways to overcome this imbalance (80, 82). A recent study also reported upregulation of nitrogen assimilation genes during infection of *C. tenuissimus* with CtenRNAV-type II (17), suggesting that this may be a conserved feature of RNA virus infection in diatoms. Furthermore, the late upregulation of ammonium, urea and amino acid transporters documented during CtenRNAV type-I infection may indicate a shift towards alternative sources of nitrogen during late-stage infection, possibly due to the availability of organic nitrogen compounds released during advanced virus infection and at the onset of host lysis.

#### Modifications in silicon metabolism during virus infection

To generate the silica-based cell wall (frustule), diatoms uptake dissolved silicic acid, Si(OH)_4,_ from the environment and transport it intracellularly to a specialized vesicle called the silica deposition vesicle (SDV) where it is polymerized into silica and assembled into a species-specific, nanoscale pattern by a protein-based scaffold (reviewed in 72). At high concentrations, Si(OH)_4_ can freely diffuse across the cell membrane, but at low concentrations, uptake is mediated through plasma-membrane bound silicon transporters (SITs; 73, 74). *C. tenussiumus* has eight predicted SIT genes (Data S6) with only one SIT (GFH46624.1) exhibiting differential expression during infection with ∼4-fold upregulation at 3 dpi and ∼4-fold downregulation at 4 dpi (Fig. 5A).

**Figure 5.**
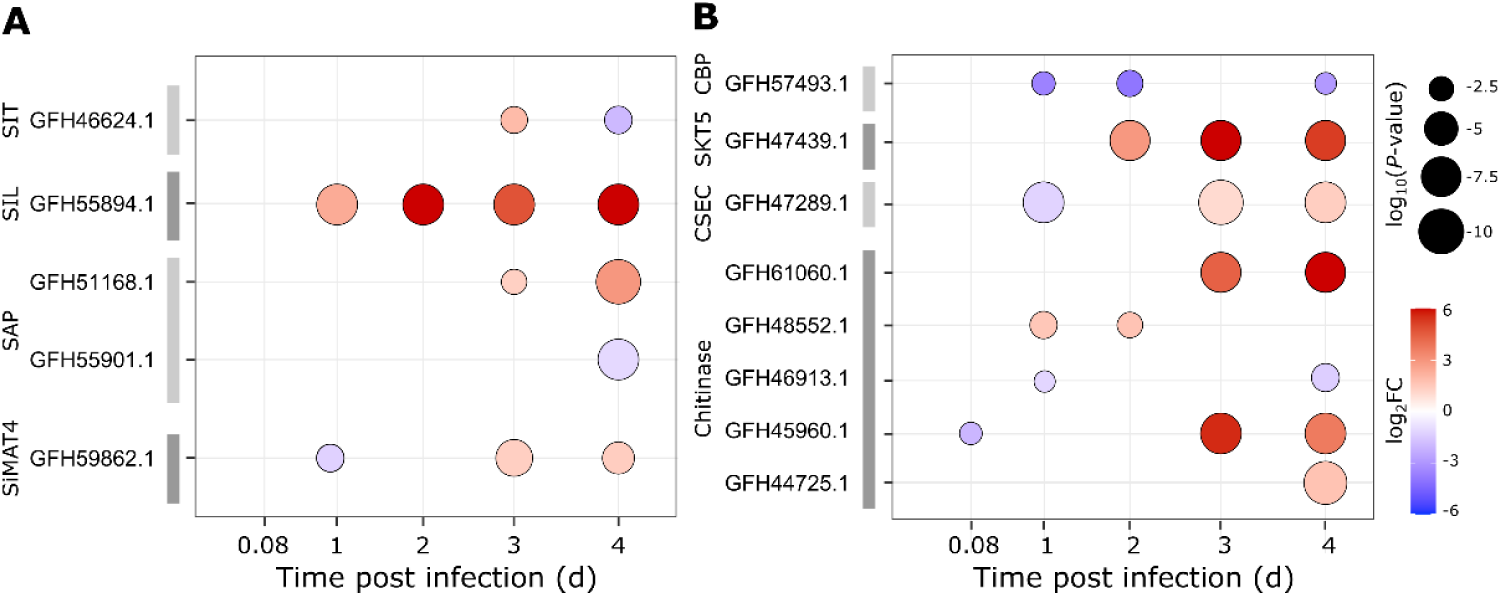
Remodeling of cell wall metabolism during CtenRNAV virus infection. Differentially expressed genes related to **A**) silicon and **B**) chitin metabolism in *C. tenuissimus* throughout the course of CtenRNAV virus infection as compared to uninfected, Ctrl cultures. Silicon metabolism genes include silicon transporters (SIT), silaffins (Sil), silacanin-1 (Sin1), silica matrix protein (SiMat), and silicalemma-associated proteins (SAP). Chitin metabolism-related genes include a chitin-binding type-4 domain-containing protein (CBP), a chitin synthase regulator (CSR, SKT5), a chitin synthase export chaperone (CSEC) and several chitinases. Symbol color depicts Fold Change (log_2_FC) and symbol size depicts adjusted *P*-values [log_10_(*P*-value); FDR] for each pairwise comparison. Only genes that were significantly differentially expressed (|log_2_FC| > 1, FDR < 0.05) are shown. Full description of gene annotations, fold change and adjusted *P*-values are available in Data S2, S3 and S6.

To further examine virus-mediated impacts on silicon metabolism, we manually annotated homologs of other known silicon metabolism-related genes (see Methods) associated with the SDV, including silaffins (Sil), silica matrix proteins (SiMat), and silicalemma-associated proteins (SAPs; Data S6). Silaffins are polycationic peptides that facilitate the polymerization of silica in the SDV (86). The family of SiMat proteins, which includes Silicanin-1 (Sin1, formerly SiMat7), are transmembrane proteins localized to the SDV that play a role in the assembly of the silica-forming organic template (87). SAPs are also associated with the SDV membrane and are thought to play a role in facilitating interactions with cytoplasmic proteins, possibly cytoskeleton-related, during pattern formation (88).

We identified 84 putative Sil genes in the *C. tenuissimus* genome (Data S6). Starting at 1 dpi, there was an increasing number of genes that were upregulated in CtenRNAV-infected cells (15, 6, 30, and 45 on 1, 2, 3, 4 dpi, respectively) with one (GFH55894.1) exhibiting substantial upregulation (∼5-640-fold) across all 4 timepoints (Fig. 5A). In contrast, only 4-9 putative silaffins were downregulated throughout infection (Data S6). Neither of the identified Sin1 homologs (GFH55092.1 and GFH43616.1) were differentially expressed at any time during CtenRNAV virus infection. Only one SiMat4 homolog was identified (GFH59862.1), which was ∼2.4-fold downregulated 1 dpi but then upregulated ∼2.6-fold 3 and 4 dpi (Fig. 5A). In addition, one of four SAP genes identified in *C. tenuissimus* (GFH51168.1; Data S6) was ∼2-8 upregulated at 3 and 4 dpi, respectively, while another SAP gene (GFH55901.1) was ∼2-fold downregulated at 4 dpi in infected cells (Fig. 5A). These data indicate that there is not a substantial shift in SIT-mediated Si uptake during CtenRNAV virus infection, but the increasing upregulation of silaffins in infected cells suggests there may be an impact on silica polymerization.

In addition to silica, chitin is one of the most abundant carbohydrates associated with the diatom cell wall (89, 90). An organic, nitrogen-containing, long-chain biopolymer of N-acetylglucosamine, chitin provides structural support for the cell wall, and in some species is extruded through pores in the frustule. In *T. pseudonana*, chitin synthases are localized near the silicified girdle bands (91) and expression increases in response to silicon limitation (92). Additionally, chitinase gene expression in *T. pseudonana* is upregulated by diverse abiotic stressors, including silicon depletion (93).

A chitin binding protein (CPB; GFH57493.1) was among the most substantially downregulated genes throughout CtenRNAV infection (12-, 13- and 8-fold at 1, 2, and 4 dpi, respectively; Fig. 5B). This was concurrent with sustained upregulation (∼8 - 80-fold) of a chitin-synthase regulator (SKT5; GFH47439.1) at 2-4 dpi and 2-3-fold upregulation of a chitin synthase export chaperone (CSEC; GFH47289.1) at 3-4 dpi compared to uninfected cells (Fig. 5B). Multiple putative chitinases were also upregulated in infected cells, increasing ∼3-fold by 1-2 dpi (GFH48552.1), ∼3-fold (GFH44725.1) by 4 dpi and ∼20-90-fold (GFH61060.1) by 3 - 4 dpi (Fig. 5B). An additional chitinase (GFH45960.1) was initially ∼4-fold downregulated at 0.08 dpi in infected cells but subsequently increased 49-fold and 13-fold by 3 and 4 dpi, respectively (Fig. 5B).

Interestingly, some viruses, including certain *Phycodnaviridae*, encode for chitinases (94) that are thought to be involved in cell wall degradation to enable the release of virus progeny (95). In diatoms, chitinase production by algicidal bacteria was shown to mechanistically degrade the cell wall in *T. pseudonana*, leading to algal lysis and death (96). Together, these data raise the possibility that the observed upregulation in chitinase during CtenRNAV virus infection may serve to permeabilize the diatom frustule, enabling virus exit from the host cell. Detailed microscopic analysis of frustules produced during infection would provide valuable insight into virus-specific impacts on silica morphogenesis and diatom cell wall integrity. Alternatively, as chitinases break down chitin into N-acetylglucosamine which can be further broken down into ammonium, the increased chitinase expression observed during CtenRNAV infection may be an additional mechanism employed by the virocell to recycle intracellular nitrogen. Chitin was previously shown to serve as an bioavailable source of nitrogen in multiple species of algae and cyanobacteria (97).

### Lipidomic remodeling in *C. tenuissimus* during CtenRNAV virus infection

#### Virus-induced shifts in membrane lipid composition

We analyzed the lipid composition in both Ctrl and CtenRNAV-infected *C. tenuissimus*, identifying 593 unique intact polar lipid and triacylglycerol (TAG, i.e. fat) molecules across all samples (Fig. 6, Data S7). Phosphatidylglycerol (PG), phosphatidylinositol (PI), and lyso-phosphatidylcholine (LPC; i.e. phosphatidylcholine, PC, with only one fatty acid) were the most abundant phospholipids in uninfected Ctrl cells, comprising ∼25%, ∼11%, and ∼8% of total non-pigment lipids, respectively, across all timepoints (Fig. S6; Data S8). Phosphatidylsulfocholine (PSC), a rare sulfur-based analog of PC (98) comprised ∼2.2% of total lipids; lyso-phosphatidylethanolamine (L-PE) comprised ∼2.3% of total lipids; and, surprisingly, intact phosphatidylcholine (PC) comprised only ∼0.9% of total lipids in Ctrl samples (Fig. S6).

**Figure 6.**
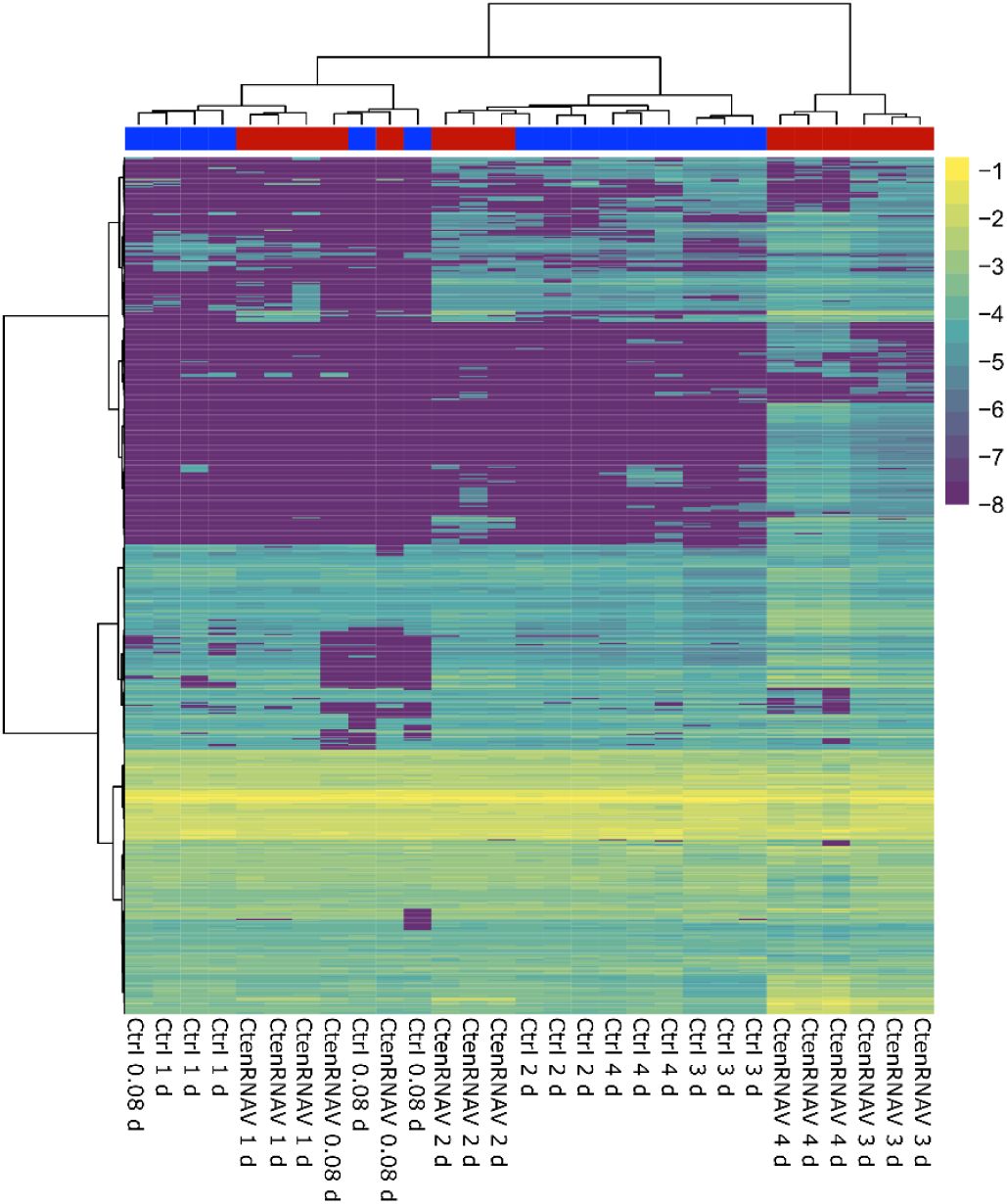
Remodeling of the *C. tenuissimus* lipidome during CtenRNAV infection. Heatmap and cluster dendrogram of 546 unique intact polar lipid and triacylglycerol molecule relative abundance in log_10_ transformed color space with cluster dendrogram of uninfected, Ctrl (blue) and CtenRNAV-infected (red) treatments (n = 3). For visual clarity, undetected lipids were annotated as 10^-8^ fraction of total lipids. See Data S7 for full list of annotated lipids.

The cellular abundance of most polar lipid classes stayed relatively similar in CtenRNAV-infected and Ctrl cells throughout infection (Fig. S7), suggesting most lipid membrane structures remained intact. However, as the infection progressed, there was a clear shift in lipid composition in the cellular molar abundance of specific lipids (Fig. 7), driving changes in the relative abundance of all lipid classes (Fig. S6), consistent with the observed transcriptomic shift in lipid metabolism-related genes (Fig. 2). Particularly notable was the abundance of phosphatidylethanolamine (PE), which was the only phospholipid class that showed consistent, significant differences between Ctrl and infected cells in per-cell abundance (Fig. 7A). In Ctrl cells, PE comprised a negligible fraction of total lipids (∼0.05%, 1.5 ± 1.1 amol cell^-1^) across all timepoints (Fig. S6, Data S8). In contrast, in CtenRNAV-infected cells, there was a ∼6-fold increase in PE within 1 dpi (Student’s t-test *P* = 0.06) that continued to increase as the infection progressed (Fig. 7A), exhibiting a ∼44-fold increase by 4 dpi compared to Ctrl cultures and accounting for ∼2% of the total lipids in infected cells (40.16 ± 11.93 amol cell^-1^; Fig. S6). All positive-strand ssRNA viruses replicate within viral replication complexes (VRC) but the type of host phospholipid is virus-specific. These data suggest that CtenRNAV may utilize PE-derived VRCs that are formed by negative curvature of the host membrane whereby the membrane invaginates away from the cytoplasm, encapsulating the viral replication proteins and protecting viral RNA synthesis from host defense mechanisms (99, 100). PE plays a role in the replication of tomato bushy stunt virus (TBSV) and the carnation Italian ringspot virus (CIRV), both belonging to the family *Tombusviridae*, as well as Nodamura virus (NoV; family *Nodaviridae*) (101). TBSV co-opts PE from the host peroxisomal membrane, while CIRV and NoV rely on mitochondrial derived PE (Zhang *et al.*, 2019).

**Figure 7.**
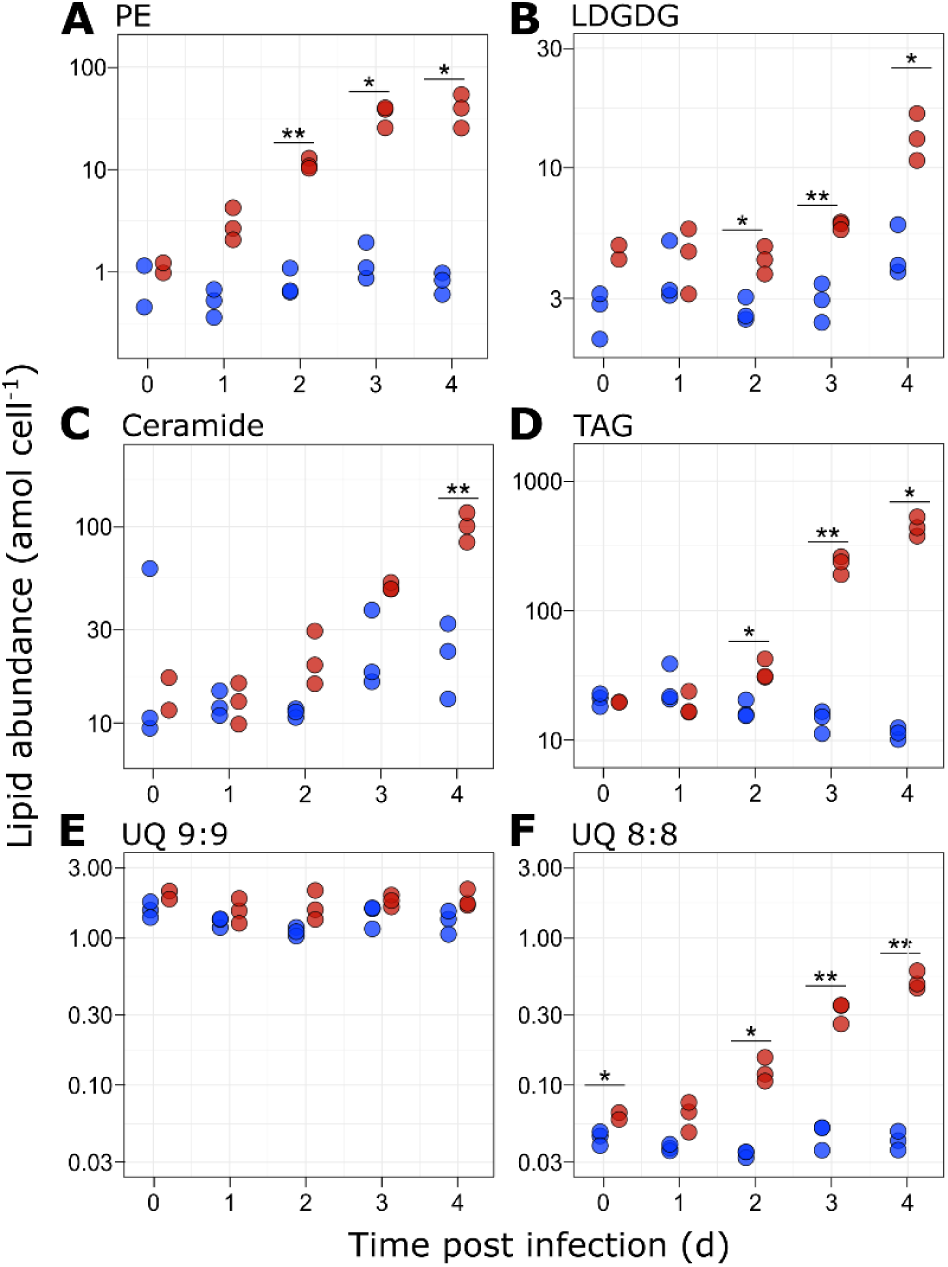
Cellular abundance of select lipids during CtenRNAV infection. Cellular abundance (amol cell^-1^) of intact membrane lipids in uninfected Ctrl (blue) and CtenRNAV-infected (red) cultures (n=3) for **A)** phosphatidylethanolamine (PE), **B)** lyso-digalactosyldiacylglycerol (LDGDG), **C)** ceramide, **D)** triacylglycerol (TAG) and electron carriers **E)** ubiquinone (UQ) 9:9 and **F)** UQ 8:8 (indicating eight and nine isoprenoid units in the molecule tails, respectively). Asterisks indicate significant differences between Ctrl and CtenRNAV-infected treatments (Student’s t-tests: \**P* < 0.05; \*\**P*< 0.01).

Shifts in FA composition were also observed in CtenRNAV-infected cells. In Ctrl cells, PE was predominantly composed of a homogeneous mix of 16 carbon (i.e. 16C) FA pairs with compounds PE 32:3, 32:4, and 32:5 (i.e. the number of carbon atoms:number of double bonds between two acyl chains; Figs. S8 and S9). In contrast, PE in infected cells was dominated by novel FA pairs that increased throughout the time course of CtenRNAV infection, driven by an increase in 18C/16C acyl chain pairs (i.e. PE 34:1, 34:2; Fig. S9). Virus-mediated shifts in FA composition were also apparent in PG and PC, despite the total abundance of these pools being similar to Ctrl cells (Figs. S7, S8 and S9). While PG and PC in Ctrl cells were comprised of paired medium and long-chain polyunsaturated FAs (e.g. PG 16:0/20:5 and PC 20:5/20:5), with minor abundance of 18C chains in CtenRNAV-infected cells (Fig. S8), PG and PC compounds were enriched with monounsaturated and di-unsaturated 18C fatty acids that increased ∼1-2 orders of magnitude throughout the time course of infection (e.g. PG 34:1, 34:2, 36:2, 36:3, and PC 34:1 and 34:2; Fig. S8). Notably, several of the novel 18C FA pairs identified in CtenRNAV-infected cells in both the PE and PG pools were below detection in uninfected, Ctrl cells (PE 34:1, 34:2, and 34:3; PG 36:2 and 36:3; Fig. S8), suggesting that these FAs may be specific to virus infection. Longer lipid FAs may contribute different membrane fluidities or curvatures that facilitate VRC formation (102). Notably, we did not detect significant abundances (>1 amol cell^-1^) of 34:1 or 34:2 glycerolipids in any other lipid class (Data S7), suggesting CtenRNAV infection may induce *de novo* monounsaturated 18C FAs lipid synthesis rather than a redistribution of existing cellular lipids.

Despite this substantial increase in select FA pairs, the most abundant PG and PC compounds in Ctrl cells remained prominent in CtenRNAV-infected cells, suggesting their main cellular functions stayed mostly intact in infected cells (Fig. S8). Previous research proposed independent synthesis pathways for PG versus PC and PE in diatoms, where PG synthesis occurs primarily in the chloroplast, while PC and PE production take place in the endoplasmic reticulum (ER; 90). Therefore, the inclusion of 18C FAs in novel PG compounds may suggest overproduction and unintentional incorporation of 18C acyl chains along another synthesis pathway.

We also observed enrichment and changes in FA composition in several mono-acyl lyso-lipid classes. Although lyso-lipids are often considered degradation products, they can also function as messenger molecules, reaction intermediates, and sources of carbon for respiration (104). In CtenRNAV-infected cells, the glycolipid, lyso-digalactosyldiacylglycerol (LDGDG) increased ∼1.6-, 2.0-, and 2.9-fold as compared to Ctrl cultures at 2, 3, and 4 dpi (Fig. 7B), though we did not detect any 18C LDGDG in either Ctrl or CtenRNAV-infected cells. In contrast, total LPC and LPE were significantly enriched in CtenRNAV-infected cells only at 4 dpi (Fig. S7; Student’s t-test *P* < 0.05); however, the individual mono-acyl compounds LPE 16:1, 17:1 and 18:1 and LPC 16:0, 16:1, and 18:1 consistently increased in abundance in infected cells compared to Ctrl cells throughout the time course of infection (Fig. S9). The lyso-glycolipid LMGDG has been identified as a precursor to TAG synthesis in the green alga, *Chlamydomonas reinhardtii* (105), and LPC and LPE are both intermediates in polyunsaturated FA synthesis in the diatom *P. tricornutum* (106), as well as phospholipid and TAG synthesis in plants (107), suggesting LDGDG, LPC, and LPC could function similarly in *C. tenuissimus*. However, lyso-lipids like LPC can be as abundant in diatoms as their diacyl counterparts (i.e. PC; 95), suggesting mono-acyl lipids may perform core cell functions beyond production and consumption as reaction intermediates or degradation products.

There was also a consistent increase in ceramide concentration in CtenRNAV-infected cells throughout the course of infection, with a ∼4.4-fold increase in infected cells compared to Ctrl by 4 dpi (Fig. 7C). Ceramide lipids have a sphingosine backbone that plays a critical role in nearly every step of the viral life cycle, including viral entry, replication, and release, across diverse host-virus lineages (109). Similar to PE, increased ceramide concentration within a lipid bilayer can induce negative membrane curvature (110), which could aid in VRC formation. Notably, in contrast with the phospholipids, we did not detect any major differences in the sphingosine/FA pair composition between CtenRNAV-infected and Ctrl cells (Fig. S8), suggesting innate *C. tenuissimus* ceramides function adequately in VRC formation or that increased ceramide production may be involved in the cellular response to virus infection (109). We did not detect glycosphingolipids, or other ceramide-based intact lipids such as phosphatidylinositol-4-phosphate, which have been shown to play significant roles during virus infection in other model systems (100, 111).

Other major components of the *C. tenuissimus* lipidome showed no significant change during viral infection (Fig. S7). The chloroplast-associated glycolipids, sulfoquinovosyldiacylglycerol (SQDG), digalactosyldiacylglycerol (DGDG), and monogalactosyldiacylglycerol (MDGD) were not significantly different between infected and Ctrl cultures (Fig. S6; Fig. S7). Additionally, we did not detect phosphatidylserine (PS) in any sample (Data S7), indicating the PS-mediated virus infection mechanisms observed in other systems (100, 112) do not play a role during CtenRNAV infection in *C. tenuissimus*.

#### Modification of lipid carbon allocation and energetic metabolism during virus infection

*C. tenuissimus* also exhibited signs of significant modulation in lipid carbon storage and metabolism during CtenRNAV infection. At 0.08 dpi, both Ctrl and infected cells exhibited similar low cellular TAG concentrations (∼20 amol cell^-1^; Fig. 7D). However, apparent by 2 dpi, TAG production in infected cells increased to 447.94 ± 77.43 amol cell^-1^ by 4 dpi, compared to only 11.38 ± 1.19 amol cell^-1^ in Ctrl cultures (Fig. 7D), and accounted for ∼25% of total lipids in CtenRNAV-infected cells (Fig. S6). Of the eight identified putative diacylglycerol acyltransferase (DGAT)-encoding genes (EC 2.3.1.20), the enzyme that catalyzes the final step in TAG biosynthesis via the acyl-CoA-dependent Kennedy pathway (113), two were downregulated (∼2-5 fold; GFH55000.1, GFH57609.1) in infected cells at 3 and 4 dpi compared to Ctrl, and one was ∼3-fold upregulated at 4 dpi (GFH50318.1; Data S2). Although DGAT characterization in diatoms is limited to a few model species, diatom genomes encode multiple, differentially localized and regulated DGAT enzymes that exhibit specific substrate affinities (i.e. for acylCoA bearing 16:0 FA versus 18:1 FA; 103). A putative phospholipid:diacylglycerol acyltransferase (PDAT; GFH59460.1), which mediates TAG biosynthesis through an acyl-CoA-independent pathway, was ∼3-fold upregulated at 1 dpi in infected cultures, followed by ∼3-fold downregulation at 3-4 dpi (Data S2), suggesting this enzyme was not involved in the TAG accumulation during late-stage CtenRNAV infection. TAG accumulation was observed in infected *G. huxleyi*, together with simultaneous up- and down-regulation of several host DGAT genes (114).

TAG accumulation is also a well-documented response to nitrogen limitation in diatoms (115–118), redirecting carbon partitioning towards lipid storage. This is consistent with the observed upregulation in nitrogen assimilation genes during CtenRNAV infection (Fig. 4). In addition, enhancing TAG production is a target in algal-based biofuel production research (113, 119, 120). We observed a ∼40-fold increase in TAG production during late CtenRNAV infection in *C. tenuissimus* (Fig. 7D), substantially greater than a ∼2-fold increase in TAG production induced by overexpression of DGAT in *P. tricornutum* and *T. pseudonana* (121–123). Although healthy *C. tenuissimus* maintains a lower constitutive TAG abundance than *P. tricornutum* and *T. pseudonana*, these data support an intriguing potential application of algal viruses in biofuel production (114, 124).

We additionally observed a shift in TAG FA composition in infected cells ( Figs. S10, S11). The most abundant TAG species in Ctrl cells contained a polyunsaturated fatty acid (e.g. TAG 46:5, 48:5, 48:6, 50:6, Fig. S10), whereas in CtenRNAV-infected cells, the most abundant TAG compounds at 4 dpi were mono-and di-unsaturated species (e.g. TAG 46:1, 48:1, 48:2, Fig. S10). This suggests there was a moderate shift in acyl-CoA substrate preferences in the TAG biosynthesis pathway during CtenRNAV infection, likely induced by changes in DGAT regulation.

The shift in phospholipid and TAG FA composition without apparent changes in glycolipid FA composition suggests enhanced lipid biosynthesis in the ER but not the plastid (103, 125). The formation of new membranes in the ER is consistent with the ER membrane being a progenitor of PE-derived VRCs in some RNA viruses (99), as well as a source of autophagosome membrane lipids in plants (126). The ER is also the primary location of lipid droplet formation in diatoms. Lipid droplets are critical energy sources for virus replication across taxonomically diverse viruses (100).

A virus-induced increase in cellular TAG and shifting FA composition may also impact diatom nutritional quality. Microalgal fatty acid composition, especially the abundance of polyunsaturated fatty acids, was linked to the reproductive success in the copepod *Temora longicornis*, and microalgal quality was shown to limit copepod reproduction in the North Sea (127). While TAG accumulation during CtenRNAV infection may augment labile carbon availability in surface ocean food webs, the apparent relative increase in saturated and monounsaturated FAs in virocells may ultimately reduce the nutritional quality of infected *C. tenuissimus,* as algae are the only producers of essential polyunsaturated fatty acids in animal and human diets (128). Moreover, the viral-induced restructuring of the lipidome may influence remineralization and particle buoyancy after host lysis due to variations in bacterial clade genetic capacities to degrade specific lipid classes (129).

Significant increases in respiration-related lipid compounds were also measured in CtenRNAV-infected cells (Fig. 7E-F). Ubiquinones (UQ) are a group of lipid soluble electron carriers in the mitochondria (130) that also function as ROS scavengers in fatty acid metabolism (131). While the cellular concentration of the primary UQ compound identified in *C. tenuissimus*, UQ 9:9 (indicating nine isoprenoid units in its tail), was not significantly different between infected and uninfected cells (Fig. 7E), there was a significant increase in a secondary quinone, UQ 8:8, in CtenRNAV-infected cells (Fig. 7F). The cellular concentration of UQ 8:8 increased ∼12-fold at 4 dpi in infected cultures compared to uninfected Ctrl cultures (Fig. 7F). While UQ 8:8 abundance was roughly an order of magnitude lower than UQ 9:9 in uninfected Ctrl cells across all timepoints, cellular UQ 8:8 and UQ 9:9 concentrations were comparable in magnitude during late stage CtenRNAV-infected cells (1.83 ± 0.26 amol_UQ 9:9_ cell^-1^ vs. 0.51 ± 0.07 amol_UQ 8:8_ cell^-1^, respectively; Fig. 7E-F). This increased ubiquinone expression suggests a significant increase in cellular oxidative phosphorylation during CtenRNAV infection, possibly to support increased host energetic demands during viral replication, or as an antioxidant response to increased ROS production in infected cells. In support of this hypothesis, we also observed increased abundance of oxidized lipids, primarily TAG, in CtenRNAV-infected cells (Fig. S12), consistent with transcriptomic indications of increased cellular stress during late-stage virus infection (Fig. 3).

### Implications of RNA virocell transformation in marine diatoms

Our findings demonstrate that CtenRNAV type-I virus infection of *C. tenuissimus* generates extensive remodeling of cellular stress, signaling and PCD-like pathways together with virus-mediated control of host resources prior to the onset of host lysis. Lipidomic remodeling and stimulation of lipid biosynthesis in infected cells implicate ER-derived PE in VRC formation, a mechanism observed in plant RNA viruses and suggestive of a strikingly conserved viral replication strategy across these evolutionary distinct red- and green-plastid host lineages (132).

The observed metabolic reprogramming during CtenRNAV virus infection in *C. tenuissimus*, including the upregulation of nitrogen assimilation pathways and pronounced TAG accumulation, is consistent with the induction of a functional nitrogen limitation response in infected diatoms. Notably, these responses occurred under nitrate-replete conditions, indicating a coordinated, virus-mediated effort to increase nitrogen supply rather than the consequence of resource depletion. Given that marine diatoms are already outstanding competitors for nitrogen (133), diatom virocells may exert an even greater competitive advantage in acquiring extracellular nitrogen within diverse phytoplankton assemblages. These findings also raise the possibility that elevated expression of nitrogen assimilation genes in natural diatom populations may signify active virus infection, rather than explicit nitrogen limitation.

Our findings indicate that RNA virus infection transforms diatom cells into metabolically distinct entities, that actively reshape nutrient cycling, carbon allocation and lipid composition prior to the onset of host lysis. The accumulation of TAGs in infected diatoms indicates a shift in carbon partitioning and supports the hypothesis that virocell transformation in diatoms may alter trophic transfer and energy flow via changes in diatom nutritional quality. Together, these findings extend the functional role of RNA viruses beyond agents of diatom mortality, suggesting that diatom virocells constitute a transient but ecological and biogeochemically consequential cell state within a broader phytoplankton community.

## Methods

### Culture conditions

*Chaetoceros tenuissimus* Meunier NIES-3715 (previously strain 2-10) and CtenRNAV type-I (family *Marnaviridae*, genus *Bacillarnavirus*; (47, 67), isolated from the coastal waters of Japan, and its associated viruses were kindly provided by Dr. Yuji Tomaru at the National Research Institute of Fisheries and Environment of Inland Sea, Japan. Host cultures were maintained in modified SWM-3 media (134) with 0.2 mM silicate, 2 mM nitrate and 2 nM Na_2_SeO_3_, at 15°C on a 12:12 light:dark cycle at ∼120 μmol photons m^−2^ s^−1^. Cell abundance was measured by flow cytometry (BD Accuri C6; 488 nm excitation, > 670 nm emission) using chlorophyll fluorescence and forward scatter. Bacterial concentrations, quantified by SYBR staining and flow cytometry, were consistently low ranging from 5 x 10^4^ – 3 x 10^5^ cells mL^-1^ (23). Cultures were infected at mid-exponential phase (∼2-5 x 10^5^ cells mL^-1^) at a multiplicity of infection (MOI) of 10. Viral lysates were generated by filtering lysed cultures through a 0.22 µm-pore sized PVDF filter and stored at 4°C.

### Biophysical measurements

Maximum photochemical quantum yield of photosystem II (photosynthetic efficiency; F_v_/F_m_) was measured using a custom-built Fluorescence Induction and Relaxation System (135), providing measurements of F_o_ (minimum) and F_m_ (maximum) fluorescence yields. Maximum efficiency of photosystem II (PSII) was calculated as F_v_/F_m_ = (F_m_ − F_o_)/F_m_.

### Virus quantification

Biomass samples for intracellular virus quantification were collected onto 1.2 µm membrane polycarbonate filters, flash frozen in liquid N_2_ and stored at -80°C until extraction. RNA was extracted using TRIzol reagent (Thermo Fisher Scientific) according to manufacturer’s instructions. Samples were treated with TURBO DNAse (Invitrogen) and RNA was quantified using a Nanodrop 1000 (Thermo Scientific). Complementary DNA (cDNA) was prepared from 800 ng total RNA using the SuperScript III First Strand Synthesis system with random hexamers (Invitrogen). Quantitative reverse-transcription PCR (qRT-PCR) was performed on a MX3000P system (Strategene) using PowerUp SYBR Green Master Mix (Applied Biosystems) in 10 μl triplicate reactions. Reactions targeted the CtenRNAV replicase polyprotein gene (gp1) with the primer pair CtenRNAV_4229F: 5′-GGCTGGTGCTTACAACGCG –3′ and CtenRNAV_4312R: 5′– CGGCATGTGCAAGTGGCT - 3′, yielding a 101 bp amplicon under the following reaction conditions: 95⁰C for 2 min, 40 cycles of 95⁰C for 30 s, 56⁰C for 30 s and 72⁰C for 45 s. Standard curves for gene-copy quantification and primer efficiencies were generated using gel-purified amplicons (PureLink Quick Gel). Reactions with no template added served as negative controls.

Extracellular virus abundance was measured in lysates generated by filtering cultures through a 0.22-µm pore-size filter to remove cellular debris. Lysates were stored at 4°C and processed within one month of collection. The concentration of infectious virus particles (infectious units mL^-1^) was measured using the most-probable-number assay (136) as described previously (32) and calculated using the EPA-MPN calculator (137).

### RNA extraction and sequencing

Biomass samples for RNA-seq were collected onto 1.2 µm polycarbonate membrane filters, resuspended in 0.2 µm-filtered seawater and pelleted by centrifugation at 4,000 x g for 10 minutes at 4°C. After removing the supernatant, pellets were flash frozen in liquid N_2_ and stored at -80° C. RNA was extracted using the Rneasy Plant Mini kit (Qiagen) and the Plant RNA Isolation Aid (Invitrogen). Library prep and sequencing were performed at Genewiz Sequencing Center (Plainfield, NJ). Libraries were constructed using the Ultra II RNA Library Prep kit (NEBNext) with Poly-A selection. Libraries were then sequenced (2 x 150 bp) on an Illumina HiSeq platform.

### Bioinformatics and statistical analysis

Poly-A/T tails and Illumina adapters were trimmed using cutadapt (138) and resulting reads shorter than 30 bp were discarded. Reads were mapped to the *C. tenuissimus* strain NIES-3715 reference genome (GenBank Accession No. GCA_021927905.1) (48) and the CtenRNAV genome (NCBI Accession No. NC_038321.1) (47) using STAR (139). Expression levels for each gene were quantified using htseq-count (140), with a concatenated gtf file of the two genomes. Differentially expressed genes were identified using DESeq2 (141) with the betaPrior, cooksCutoff and independentFiltering parameters set to False. Raw P-values were adjusted for multiple testing using the procedure of Benjamini and Hochberg. Pipeline was run using snakemake (142). Only genes that had a count of at least 30 reads in at least one sample were selected for downstream analysis. Differentially expressed genes (DEGs) were defined as genes with a log_2_ fold change (|log_2_FC|) > 1 and adjusted *P*-value (FDR) < 0.05. DEGs occurring in at least two comparisons were clustered using hierarchical clustering with Ward linkage (143, 144). Significantly enriched GO terms within each cluster were identified using blast2go (145) and a hypergeometric enrichment test with *P* ≤ 0.05 as the threshold for significance.

### C. tenuissimus ortholog identification and functional annotation

Within the recently published *C. tenuissimus* genome (GenBank Accession No. GCA_021927905.1) (48), orthologous proteins were identified using reciprocal blast (BLASTP; E-value threshold < 10^-5^) against National Center for Biotechnology Information (NCBI) reference genome sequences and protein datasets for *Phaeodactylum tricornutum* (GCF_000150955.2), *Thalassiosira pseudonana* (GCF_000149405.2), and *Saccharomyces cerevisiae* (GCF_000146045.2) (146, 147). An additional reciprocal blast (BLASTP) was performed against the UniProt50 database (49) to enhance annotation coverage and accuracy. For UniProt50 matches, each identifier represents a cluster of proteins sharing ≥ 50% sequence identity, from which one representative protein was randomly selected for annotation purposes. Definitive orthology and functional annotation for each putative protein sequence was retained only if the initial protein sequence was recovered in the reciprocal blast. Additional functional annotation was performed using the KEGG Orthology database through KofamKOALA (148) and PANNZER (149), with Gene Ontology (GO) predictions filtered using a Positive Predictive Value (PPV) threshold of ≥ 0.5 to ensure annotation reliability.

Silaffin-like genes were identified in the *C. tenuissimus* genome, as described previously (150, 151), using a sliding window to search translated nucleotide sequences containing a 100-2000 long segment with ≥10% lysine and ≥18% serine residues. Proteins matching the selection criteria were further screened for the presence of a signal peptide using SignalP V6.0 software with a likelihood cutoff of 0.9. SiMAT-like genes were identified by the defined presence of at least 3 KxxK motifs (where x is glycine, serine, or alanine), 1 RxL motif, and 10-12% proline residues (152). The sequence of Silicanin-1 (Sin1) from *T. pseudonana* (Tp) was used as a query against the *C. tenuissimus* genome using BLASTP. Using the criteria defined by Kotzsch et al. (87), hits with an e-value < -50 and a RxL motif were retained. Silicalemma-associated proteins (SAPs) were identified by BLASTP using TpSAP1 and TpSAP2 as queries with an initial e-value cutoff of 10. Those with a serine rich region (defined as having at least 50% serine residues across 10-40 amino acids) and transmembrane domains upstream of the conserved cytoplasmic domain (88) were considered SAP-like proteins.

### Lipid analysis

For lipid analysis, 20 mL aliquots of culture replicates were filtered onto 0.2 µm Durapore filters (Millipore, USA), flash frozen in liquid nitrogen, and stored at -80°C until analysis. Samples were extracted using a modified Bligh-and-Dyer procedure (153), according to Popendorf et al. (154), during which a suite of deuterated lipid standards was added to the extraction vial for individual compound class recovery correction. Aliquots of the lipid extracts were dried down and resuspended in a 7:3 (v/v) mixture of acetonitrile:isopropanol (155), then analyzed via high-performance liquid chromatography tandem high-resolution accurate-mass mass spectrometry (156). Subsequently, mass spectral data were retention time corrected and annotated using the LOBSTAHS package (157) in the R-Language, which uses the xcms (158) and CAMERA (159) packages. Lipid assignments were verified using a combination of exact mass, ms2 fragmentation, feature matching in positive and negative ionization modes, and retention time patterns based on fatty acid carbon numbers and double bonds (156, 160). Lipid mass spectral assignments were then quantified using ionization coefficient equations calculated from external standards curves for each major lipid class, except where none were available; in those cases, we proceeded as follows: for phosphatidylsulfocholine (PSC), we used the ionization coefficient for phosphatidylethanolamine as a “best-matched” standard based on analogous headgroup structure (161), and for ceramides, we used the correction equation for a glycosylceramide. In the case of the lyso-lipids, LPG, LMGDG, and LDGDG, where commercial standards were also not available, approximate quantification coefficients were calculated by multiplying the ionization coefficients for their intact counterparts (i.e. PG, MGDG, and DGDG) by the mean ratio of the ionization coefficients of LPE and LPC to PE and PC, under the assumption that the ionization efficiency relationship between a lyso-lipid and its corresponding intact diacyl lipids stays roughly consistent across lipid classes. All pigments and quinones were corrected using an external standard curve of chlorophyll a.

### Data visualization and analysis

Statistical analyses were performed in the open source platform R, version 4.1.2 “Bird Hippie”. Data visualization and plotting were produced using the R packages ggplot2 (162), pheatmap (163) and patchwork (164).

### Data Availability

Raw sequencing data will be available upon publication at the National Center for Biotechnology Information Sequence Read Archive under NCBI’s SRA under Bioproject accession no. PRJNA1434310. Raw mass spectral lipid data files are available through the MetaboLights database. All data generated or analyzed during the current study are included in this published article and its supplementary information files.

## Author Contributions

C.F.K. and K.T. conceived the project. C.F.K., D.L. and K.T. wrote the manuscript; C.F.K. and K.T. conducted the infection experiments and sample collection; C.F.K. performed RNA extractions and qRT-PCR analysis; H.W.B. and E.Z. performed bioinformatic analysis; C.F.K. and L.B. conducted transcriptomics analysis; J.M.T and K.T. annotated and analyzed silicon metabolism-related genes; D.L, H.F. and B.A.S.V.M conducted lipid extractions and measurements; D.L. performed lipid analysis; All authors provided feedback and approved the final manuscript.

## Acknowledgements

This work was supported by an Institute of Earth, Ocean and Atmospheric Sciences (EOAS) Rutgers Postdoctoral Fellowship (C.F.K.) a Simons Foundation Postdoctoral Fellowship SF-548156 (C.F.K.), an Israel Science Foundation Grant ISF-1355/22 (C.F.K) and an Alon Fellowship (C.F.K.). Support was also provided by grants from the National Science Foundation OCE-2049386 and OIA-2021032 (K.T.). B.A.S.V.M. and D.P.L were supported by grants from the Simons Foundation (721229) and the National Science Foundation (OCE-2020878, OCE-2022597, and OPP-2026045). We thank Yuji Tomaru (National Research Institute of Fisheries and Environment of Inland Sea, Japan) for providing *C. tenuissimus* and CtenRNAV and Amir Szitenberg at the Mantoux Bioinformatics Institute (Nancy and Stephen Grand Israel National Center for Personalized Medicine, Weizmann Institute of Science) for assistance with bioinformatics analysis.

## Competing interests

The authors declare no competing interests.

## Notes

### Competing Interest Statement

The authors have declared no competing interest.

## References

1. C. B. Field, M. J. Behrenfeld, J. T. Randerson, P. Falkowski, Primary production of the biosphere: integrating terrestrial and oceanic components. Science. 281, 237–240 (1998).

2. Ø. Bergh, K. Y. Børsheim, G. Bratbak, M. Heldal, High abundance of viruses found in aquatic environments. Nature 340, 467–468 (1989).

3. S. W. Wilhelm, C. A. Suttle, Viruses and nutrient cycles in the sea: viruses play critical roles in the structure and function of aquatic food webs. Bioscience 49, 781–788 (1999).

4. C. A. Suttle, Marine viruses—major players in the global ecosystem. Nat. Rev. Microbiol. 5, 801–812 (2007).

5. C. P. D. Brussaard, Viral control of phytoplankton populations—a review 1. J. Eukaryot. Microbiol. 51, 125–138 (2004).

6. A. Vardi, et al., Host-virus dynamics and subcellular controls of cell fate in a natural coccolithophore population. Proc. Natl. Acad. Sci. 109, 19327–19332 (2012).

7. T. E. G. Biggs, J. Huisman, C. P. D. Brussaard, Viral lysis modifies seasonal phytoplankton dynamics and carbon flow in the Southern Ocean. ISME J. 15, 3615–3622 (2021).

8. W. H. Wilson, et al., Isolation of viruses responsible for the demise of an *Emiliania huxleyi* bloom in the English Channel. J. Mar. Biol. Assoc. United Kingdom 82, 369–377 (2002).

9. K. D. A. Mojica, C. P. D. Brussaard, Marine Viruses and Their Role in Marine Ecosystems and Carbon Cycling. Ann. Rev. Mar. Sci. 18 (2025).

10. C. P. Laber, et al., Coccolithovirus facilitation of carbon export in the North Atlantic. Nat. Microbiol. 3, 537–547 (2018).

11. L. Guidi, et al., Plankton networks driving carbon export in the oligotrophic ocean. Nature 532, 465–470 (2016).

12. D. Lindell, et al., Genome-wide expression dynamics of a marine virus and host reveal features of co-evolution. Nature 449, 83–86 (2007).

13. C. Evans, D. W. Pond, W. H. Wilson, Changes in *Emiliania huxleyi* fatty acid profiles during infection with E. huxleyi virus 86: physiological and ecological implications. Aquat. Microb. Ecol. 55, 219–228 (2009).

14. S. Rosenwasser, C. Ziv, S. G. Van Creveld, A. Vardi, Virocell metabolism: metabolic innovations during host-virus interactions in the ocean. Trends Microbiol. 24, 821–832 (2016).

15. M. Moniruzzaman, E. R. Gann, S. W. Wilhelm, Infection by a giant virus (AaV) induces widespread physiological reprogramming in *Aureococcus anophagefferens* CCMP1984--a harmful bloom algae. Front. Microbiol. 9, 752 (2018).

16. A. E. Zimmerman, et al., Metabolic and biogeochemical consequences of viral infection in aquatic ecosystems. Nat. Rev. Microbiol. (2020).

17. Y. Hongo, Y. Tomaru, Transcriptomic insights into host transcriptional manipulation by ssDNA and ssRNA viruses in the marine planktonic diatom Chaetoceros tenuissimus. Virus Res. 199605 (2025).

18. P. Forterre, The virocell concept and environmental microbiology. ISME J. 7, 233–236 (2013).

19. J. R. Brum, M. B. Sullivan, Rising to the challenge: accelerated pace of discovery transforms marine virology. Nat. Rev. Microbiol. 13, 147–159 (2015).

20. C. Lønborg, M. Middelboe, C. P. D. Brussaard, Viral lysis of *Micromonas pusilla*: impacts on dissolved organic matter production and composition. Biogeochemistry 116, 231–240 (2013).

21. C. Kuhlisch, et al., Viral infection of algal blooms leaves a unique metabolic footprint on the dissolved organic matter in the ocean. Sci. Adv. 7, eabf4680 (2021).

22. X. Ma, M. L. Coleman, J. R. Waldbauer, Distinct molecular signatures in dissolved organic matter produced by viral lysis of marine cyanobacteria. Environ. Microbiol. 20, 3001–3011 (2018).

23. C. F. Kranzler, et al., Taxonomically distinct diatom viruses differentially impact microbial processing of organic matter. Sci. Adv. 11, eadq5439 (2025).

24. Y. I. Wolf, et al., Origins and evolution of the global RNA virome. MBio 9, 10–1128 (2018).

25. G. F. Steward, et al., Are we missing half of the viruses in the ocean? ISME J. 7, 672–679 (2013).

26. J. A. Miranda, A. I. Culley, C. R. Schvarcz, G. F. Steward, RNA viruses as major contributors to Antarctic virioplankton. Environ. Microbiol. 18, 3714–3727 (2016).

27. Y. I. Wolf, et al., Doubling of the known set of RNA viruses by metagenomic analysis of an aquatic virome. Nat. Microbiol. 5, 1262–1270 (2020).

28. A. Culley, New insight into the RNA aquatic virosphere via viromics. Virus Res. 244, 84–89 (2018).

29. G. Dominguez-Huerta, et al., Diversity and ecological footprint of Global Ocean RNA viruses. Science. 376, 1202–1208 (2022).

30. L. Arsenieff, K. Kimura, C. F. Kranzler, A.-C. Baudoux, K. Thamatrakoln, “Diatom viruses” in The Molecular Life of Diatoms, (Springer, 2022), pp. 713–740.

31. K. Nagasaki, Dinoflagellates, diatoms, and their viruses. J. Microbiol. 46, 235–243 (2008).

32. C. F. Kranzler, et al., Silicon limitation facilitates virus infection and mortality of marine diatoms. Nat. Microbiol. 4, 1790–1797 (2019).

33. C. F. Kranzler, et al., Impaired viral infection and reduced mortality of diatoms in iron-limited oceanic regions. Nat. Geosci. 14, 231–237 (2021).

34. Y. Tomaru, N. Fujii, S. Oda, K. Toyoda, K. Nagasaki, Dynamics of diatom viruses on the western coast of Japan. Aquat. Microb. Ecol. 63, 223–230 (2011).

35. L. Arsenieff, et al., First viruses infecting the marine diatom *Guinardia delicatula*. Front. Microbiol. 9, 3235 (2019).

36. Y. Tomaru, K. Toyoda, K. Kimura, Occurrence of the planktonic bloom-forming marine diatom *Chaetoceros tenuissimus* Meunier and its infectious viruses in western Japan. Hydrobiologia 805, 221–230 (2018).

37. S. Malviya, et al., Insights into global diatom distribution and diversity in the world’s ocean. Proc. Natl. Acad. Sci. 113, E1516--E1525 (2016).

38. J. J. Pierella Karlusich, et al., Patterns and drivers of diatom diversity and abundance in the global ocean. Nat. Commun. 16, 3452 (2025).

39. D. M. Nelson, P. Tréguer, M. A. Brzezinski, A. Leynaert, B. Quéguiner, Production and dissolution of biogenic silica in the ocean: revised global estimates, comparison with regional data and relationship to biogenic sedimentation. Global Biogeochem. Cycles 9, 359–372 (1995).

40. X. Jin, N. Gruber, J. P. Dunne, J. L. Sarmiento, R. A. Armstrong, Diagnosing the contribution of phytoplankton functional groups to the production and export of particulate organic carbon, CaCO3, and opal from global nutrient and alkalinity distributions. Global Biogeochem. Cycles 20 (2006).

41. V. Smetacek, et al., Deep carbon export from a Southern Ocean iron-fertilized diatom bloom. Nature 487, 313–319 (2012).

42. S. Agusti, et al., Ubiquitous healthy diatoms in the deep sea confirm deep carbon injection by the biological pump. Nat. Commun. 6, 7608 (2015).

43. D. M. Karl, M. J. Church, J. E. Dore, R. M. Letelier, C. Mahaffey, Predictable and efficient carbon sequestration in the North Pacific Ocean supported by symbiotic nitrogen fixation. Proc. Natl. Acad. Sci. 109, 1842–1849 (2012).

44. B. R. Edwards, K. Thamatrakoln, H. F. Fredricks, K. D. Bidle, B. A. S. Van Mooy, Viral Infection Leads to a Unique Suite of Allelopathic Chemical Signals in Three Diatom Host--Virus Pairs. Mar. Drugs 22, 228 (2024).

45. M. Walde, C. Camplong, C. de Vargas, A.-C. Baudoux, N. Simon, Viral infection impacts the 3D subcellular structure of the abundant marine diatom *Guinardia delicatula*. Front. Mar. Sci. 9, 1034235 (2023).

46. A. Pelusi, et al., Virus-induced spore formation as a defense mechanism in marine diatoms. New Phytol. 229, 2251–2259 (2021).

47. Y. Shirai, et al., Isolation and characterization of a single-stranded RNA virus infecting the marine planktonic diatom *Chaetoceros tenuissimus* Meunier. Appl. Environ. Microbiol. 74, 4022–4027 (2008).

48. Y. Hongo, et al., The genome of the diatom *Chaetoceros tenuissimus* carries an ancient integrated fragment of an extant virus. Sci. Rep. 11, 22877 (2021).

49. T. U. Consortium, UniProt: the Universal Protein Knowledgebase in 2023. Nucleic Acids Res. 51, D523–D531 (2022).

50. K. Mäkinen, S. De, The significance of methionine cycle enzymes in plant virus infections. Curr. Opin. Plant Biol. 50, 67–75 (2019).

51. K. E. Helliwell, et al., A novel Ca^2+^ signaling pathway coordinates environmental phosphorus sensing and nitrogen metabolism in marine diatoms. Curr. Biol. 31, 978–989 (2021).

52. K. E. Helliwell, et al., Spatiotemporal patterns of intracellular Ca^2+^ signalling govern hypo-osmotic stress resilience in marine diatoms. New Phytol. 230, 155–170 (2021).

53. A. Falciatore, M. R. d’Alcalà, P. Croot, C. Bowler, Perception of environmental signals by a marine diatom. Science. 288, 2363–2366 (2000).

54. S. van Creveld, A. Mizrachi, A. Vardi, “An ocean of signals: intracellular and extracellular signaling in diatoms” in The Molecular Life of Diatoms, (Springer, 2022), pp. 641–678.

55. A. Vardi, et al., A stress surveillance system based on calcium and nitric oxide in marine diatoms. PLoS Biol. 4, e60 (2006).

56. K. D. Bidle, Programmed cell death in unicellular phytoplankton. Curr. Biol. 26, R594–R607 (2016).

57. Y. Zhou, T. K. Frey, J. J. Yang, Viral calciomics: interplays between Ca^2+^ and virus. Cell Calcium 46, 1–17 (2009).

58. M. Campanella, et al., The coxsackievirus 2B protein suppresses apoptotic host cell responses by manipulating intracellular Ca^2+^ homeostasis. J. Biol. Chem. 279, 18440–18450 (2004).

59. B. Chen, M. E. Feder, L. Kang, Evolution of heat-shock protein expression underlying adaptive responses to environmental stress. Mol. Ecol. 27, 3040–3054 (2018).

60. K. Richter, M. Haslbeck, J. Buchner, The heat shock response: life on the verge of death. Mol. Cell 40, 253–266 (2010).

61. H.-P. Dong, et al., High light stress triggers distinct proteomic responses in the marine diatom *Thalassiosira pseudonana*. BMC Genomics 17, 1–14 (2016).

62. H. Wang, et al., Responses of marine diatom *Skeletonema marinoi* to nutrient deficiency: programmed cell death. Appl. Environ. Microbiol. 86, e02460--19 (2020).

63. A. E. Allen, et al., Whole-cell response of the pennate diatom *Phaeodactylum tricornutum* to iron starvation. Proc. Natl. Acad. Sci. 105, 10438–10443 (2008).

64. C. Lauritano, I. Orefice, G. Procaccini, G. Romano, A. Ianora, Key genes as stress indicators in the ubiquitous diatom Skeletonema marinoi. BMC Genomics 16, 1–11 (2015).

65. A. Lubkowska, W. Pluta, A. Strońska, A. Lalko, Role of heat shock proteins (HSP70 and HSP90) in viral infection. Int. J. Mol. Sci. 22, 9366 (2021).

66. K. Kimura, Y. Tomaru, Discovery of two novel viruses expands the diversity of single-stranded DNA and single-stranded RNA viruses infecting a cosmopolitan marine diatom. Appl. Environ. Microbiol. 81, 1120–1131 (2015).

67. M. Vlok, A. S. Lang, C. A. Suttle, Application of a sequence-based taxonomic classification method to uncultured and unclassified marine single-stranded RNA viruses in the order *Picornavirales*. Virus Evol. 5, vez056 (2019).

68. K. D. Bidle, L. Haramaty, J. e Ramos, P. Falkowski, Viral activation and recruitment of metacaspases in the unicellular coccolithophore, Emiliania huxleyi. Proc. Natl. Acad. Sci. 104, 6049–6054 (2007).

69. K. D. Bidle, S. J. Bender, Iron starvation and culture age activate metacaspases and programmed cell death in the marine diatom *Thalassiosira pseudonana*. Eukaryot. Cell 7, 223–236 (2008).

70. D. Spungin, K. D. Bidle, I. Berman-Frank, Metacaspase involvement in programmed cell death of the marine cyanobacterium *Trichodesmium*. Environ. Microbiol. 21, 667–681 (2019).

71. S. van Creveld, et al., Biochemical characterization of a novel redox-regulated metacaspase in a marine diatom. Front. Microbiol. 12, 688199 (2021).

72. A. Mizrachi, et al., Cathepsin X is a conserved cell death protein involved in algal response to environmental stress. Curr. Biol. 35, 2240–2255 (2025).

73. T. J. Browning, C. M. Moore, Global analysis of ocean phytoplankton nutrient limitation reveals high prevalence of co-limitation. Nat. Commun. 14, 5014 (2023).

74. S. R. Smith, et al., Evolution and regulation of nitrogen flux through compartmentalized metabolic networks in a marine diatom. Nat. Commun. 10, 4552 (2019).

75. R. H. Lampe, et al., Divergent gene expression among phytoplankton taxa in response to upwelling. Environ. Microbiol. 20, 3069–3082 (2018).

76. L. Alipanah, J. Rohloff, P. Winge, A. M. Bones, T. Brembu, Whole-cell response to nitrogen deprivation in the diatom *Phaeodactylum tricornutum*. J. Exp. Bot. 66, 6281–6296 (2015).

77. S. J. Bender, C. A. Durkin, C. T. Berthiaume, R. L. Morales, E. V. Armbrust, Transcriptional responses of three model diatoms to nitrate limitation of growth. Front. Mar. Sci. 1, 3 (2014).

78. A. Monier, et al., Host-derived viral transporter protein for nitrogen uptake in infected marine phytoplankton. Proc. Natl. Acad. Sci. 114, E7489--E7498 (2017).

79. Q. Zeng, S. W. Chisholm, Marine viruses exploit their host’s two-component regulatory system in response to resource limitation. Curr. Biol. 22, 124–128 (2012).

80. L. F. Jover, T. C. Effler, A. Buchan, S. W. Wilhelm, J. S. Weitz, The elemental composition of virus particles: implications for marine biogeochemical cycles. Nat. Rev. Microbiol. 12, 519–528 (2014).

81. Z. V Finkel, et al., Phytoplankton in a changing world: cell size and elemental stoichiometry. J. Plankton Res. 32, 119–137 (2010).

82. D. Muratore, et al., Microbial and viral genome and proteome nitrogen demand varies across multiple spatial scales within a marine oxygen minimum zone. Msystems 8, e01095--22 (2023).

83. M. Hildebrand, S. J. L. Lerch, R. P. Shrestha, Understanding diatom cell wall silicification—moving forward. Front. Mar. Sci. 5, 125 (2018).

84. M. Hildebrand, B. E. Volcani, W. Gassmann, J. I. Schroeder, A gene family of silicon transporters. Nature 385, 688–689 (1997).

85. K. Thamatrakoln, M. Hildebrand, Silicon uptake in diatoms revisited: a model for saturable and nonsaturable uptake kinetics and the role of silicon transporters. Plant Physiol. 146, 1397–1407 (2008).

86. N. Kroger, S. Lorenz, E. Brunner, M. Sumper, Self-assembly of highly phosphorylated silaffins and their function in biosilica morphogenesis. Science (80-.). 298, 584–586 (2002).

87. A. Kotzsch, et al., Silicanin-1 is a conserved diatom membrane protein involved in silica biomineralization. BMC Biol. 15, 65 (2017).

88. B. Tesson, S. J. L. Lerch, M. Hildebrand, Characterization of a new protein family associated with the silica deposition vesicle membrane enables genetic manipulation of diatom silica. Sci. Rep. 7, 13457 (2017).

89. B. Tesson, et al., Contribution of multi-nuclear solid state NMR to the characterization of the *Thalassiosira pseudonana* diatom cell wall. Anal. Bioanal. Chem. 390, 1889–1898 (2008).

90. J. McLachlan, A. G. McInnes, M. Falk, Studies on the chitan (chitin: Poly-N-acetylglucosamine) fibers of the diatom *Thalassiosira fluviatilis* hustedt: I production and isolation of chitan fibers. Can. J. Bot. 43, 707–713 (1965).

91. M. Wustmann, N. Poulsen, N. Kröger, K.-H. van Pée, Chitin synthase localization in the diatom *Thalassiosira pseudonana*. BMC Mater. 2, 1–7 (2020).

92. C. A. Durkin, T. Mock, E. V. Armbrust, Chitin in diatoms and its association with the cell wall. Eukaryot. Cell 8, 1038–1050 (2009).

93. H. Cheng, Z. Shao, C. Lu, D. Duan, Genome-wide identification of chitinase genes in Thalassiosira pseudonana and analysis of their expression under abiotic stresses. BMC Plant Biol. 21, 87 (2021).

94. E. G. de Oliveira, C. A. da C. Filho, R. A. L. Rodrigues, An overview of viral chitinases: General properties and biotechnological potential. Exp. Biol. Med. 248, 2053–2061 (2023).

95. L. Sun, B. Adams, J. R. Gurnon, Y. Ye, J. L. Van Etten, Characterization of two chitinase genes and one chitosanase gene encoded by Chlorella virus PBCV-1. Virology 263, 376–387 (1999).

96. Y. Li, et al., Chitinase producing bacteria with direct algicidal activity on marine diatoms. Sci. Rep. 6, 21984 (2016).

97. C. E. Blank, N. W. Hinman, Cyanobacterial and algal growth on chitin as a source of nitrogen; ecological, evolutionary, and biotechnological implications. Algal Res. 15, 152–163 (2016).

98. P. Bisseret, et al., Occurrence of phosphatidylsulfocholine, the sulfonium analog of phosphatidylcholine in some diatoms and algae. Biochim. Biophys. Acta (BBA)-Lipids Lipid Metab. 796, 320–327 (1984).

99. Z. Zhang, et al., Host lipids in positive-strand RNA virus genome replication. Front. Microbiol. 10, 286 (2019).

100. E. Ketter, G. Randall, Virus impact on lipids and membranes. Annu. Rev. Virol. 6, 319–340 (2019).

101. K. Xu, P. D. Nagy, RNA virus replication depends on enrichment of phosphatidylethanolamine at replication sites in subcellular membranes. Proc. Natl. Acad. Sci. 112, E1782--E1791 (2015).

102. W. A. Lathram, et al., Dissecting the biophysical mechanisms of oleate hydratase association with membranes. Front. Mol. Biosci. 11, 1504373 (2025).

103. T. Tanaka, K. Yoneda, Y. Maeda, “Lipid metabolism in diatoms” in The Molecular Life of Diatoms, (Springer, 2022), pp. 493–527.

104. F. Kong, I. T. Romero, J. Warakanont, Y. Li-Beisson, Lipid catabolism in microalgae. New Phytol. 218, 1340–1348 (2018).

105. X. Li, et al., A galactoglycerolipid lipase is required for triacylglycerol accumulation and survival following nitrogen deprivation in *Chlamydomonas reinhardtii*. Plant Cell 24, 4670–4686 (2012).

106. K. Jasieniecka-Gazarkiewicz, A. Połońska, Y. Gong, A. Banaś, Acyl-CoA: lysophosphatidylcholine acyltransferase from diatom *P. Tricornutum* efficiently remodels phosphatidylcholine containing polyunsaturated fatty acids. Sci. Rep. 14, 30970 (2024).

107. P. D. Bates, Understanding the control of acyl flux through the lipid metabolic network of plant oil biosynthesis. Biochim. Biophys. Acta (BBA)-Molecular Cell Biol. Lipids 1861, 1214–1225 (2016).

108. K. Zhang, J. Li, J. Cheng, S. Lin, Alkaline phosphatase PhoD mutation induces fatty acid and long-chain polyunsaturated fatty acid (LC-PUFA)-bound phospholipid production in the model diatom *Phaeodactylum tricornutum*. Mar. Drugs 21, 560 (2023).

109. N. Beckmann, K. A. Becker, Ceramide and related molecules in viral infections. Int. J. Mol. Sci. 22, 5676 (2021).

110. A. Draeger, E. B. Babiychuk, Ceramide in plasma membrane repair. Sphingolipids Dis. 341–353 (2013).

111. A. Vardi, et al., Viral glycosphingolipids induce lytic infection and cell death in marine phytoplankton. Science. 326, 861–865 (2009).

112. A. Amara, J. Mercer, Viral apoptotic mimicry. Nat. Rev. Microbiol. 13, 461–469 (2015).

113. Y. Xu, et al., Properties and biotechnological applications of acyl-CoA: diacylglycerol acyltransferase and phospholipid: diacylglycerol acyltransferase from terrestrial plants and microalgae. Lipids 53, 663–688 (2018).

114. S. Malitsky, et al., Viral infection of the marine alga *Emiliania huxleyi* triggers lipidome remodeling and induces the production of highly saturated triacylglycerol. New Phytol. 210, 88–96 (2016).

115. Z.-K. Yang, et al., Molecular and cellular mechanisms of neutral lipid accumulation in diatom following nitrogen deprivation. Biotechnol. Biofuels 6, 67 (2013).

116. O. Levitan, et al., Remodeling of intermediate metabolism in the diatom *Phaeodactylum tricornutum* under nitrogen stress. Proc. Natl. Acad. Sci. 112, 412–417 (2015).

117. G. d’Ippolito, et al., Potential of lipid metabolism in marine diatoms for biofuel production. Biotechnol. Biofuels 8, 28 (2015).

118. B. Huang, et al., Nitrogen and phosphorus limitations induce carbon partitioning and membrane lipid remodelling in the marine diatom *Phaeodactylum tricornutum*. Eur. J. Phycol. 54, 342–358 (2019).

119. B. Liu, C. Benning, Lipid metabolism in microalgae distinguishes itself. Curr. Opin. Biotechnol. 24, 300–309 (2013).

120. Y. Li-Beisson, J. J. Thelen, E. Fedosejevs, J. L. Harwood, The lipid biochemistry of eukaryotic algae. Prog. Lipid Res. 74, 31–68 (2019).

121. Y. Zhang, Y. Pan, W. Ding, H. Hu, J. Liu, Lipid production is more than doubled by manipulating a diacylglycerol acyltransferase in algae. GCB Bioenergy 13, 185–200 (2021).

122. J. Dinamarca, O. Levitan, G. K. Kumaraswamy, D. S. Lun, P. G. Falkowski, Overexpression of a diacylglycerol acyltransferase gene in *Phaeodactylum tricornutum* directs carbon towards lipid biosynthesis. J. Phycol. 53, 405–414 (2017).

123. K. Manandhar-Shrestha, M. Hildebrand, Characterization and manipulation of a DGAT2 from the diatom *Thalassiosira pseudonana*: improved TAG accumulation without detriment to growth, and implications for chloroplast TAG accumulation. Algal Res. 12, 239–248 (2015).

124. A. M. Lopez, Y. Choi, Z. Zhou, Enhanced biomimetic algal lipid enrichment for improved biofuel production driven by non-stress viral lysis. Bioresour. Technol. 133128 (2025).

125. O. Sayanova, et al., Modulation of lipid biosynthesis by stress in diatoms. Philos. Trans. R. Soc. B Biol. Sci. 372 (2017).

126. J. Soto-Burgos, X. Zhuang, L. Jiang, D. C. Bassham, Dynamics of autophagosome formation. Plant Physiol. 176, 219–229 (2018).

127. K. E. Arendt, S. H. Jónasdóttir, P. J. Hansen, S. Gärtner, Effects of dietary fatty acids on the reproductive success of the calanoid copepod *Temora longicornis*. Mar. Biol. 146, 513–530 (2005).

128. J. Dalsgaard, M. S. John, G. Kattner, D. Müller-Navarra, W. Hagen, Fatty acid trophic markers in the pelagic marine environment. (2003).

129. L. Behrendt, et al., Microbial dietary preference and interactions affect the export of lipids to the deep ocean. Science. 385, eaab2661 (2024).

130. B. Nowicka, J. Kruk, Occurrence, biosynthesis and function of isoprenoid quinones. Biochim. Biophys. Acta (BBA)-Bioenergetics 1797, 1587–1605 (2010).

131. A. Maroz, R. F. Anderson, R. A. J. Smith, M. P. Murphy, Reactivity of ubiquinone and ubiquinol with superoxide and the hydroperoxyl radical: implications for in vivo antioxidant activity. Free Radic. Biol. Med. 46, 105–109 (2009).

132. P. G. Falkowski, et al., The evolution of modern eukaryotic phytoplankton. Science. 305, 354–360 (2004).

133. S. E. Fawcett, B. B. Ward, Phytoplankton succession and nitrogen utilization during the development of an upwelling bloom. Mar. Ecol. Prog. Ser. 428, 13–31 (2011).

134. L. Chen, T. Edelstein, J. McLachlan, Bonnemaisonia hamifera hariot in nature and in culture. J. Phycol. 5, 211–220 (1969).

135. M. Y. Gorbunov, P. G. Falkowski, Fluorescence induction and relaxation (FIRe) technique and instrumentation for monitoring photosynthetic processes and primary production in aquatic ecosystems. Photosynth. Fundam. Asp. to Glob. Perspect. 1029–1031 (2004).

136. P. F. Kemp, J. J. Cole, B. F. Sherr, E. B. Sherr, Handbook of methods in aquatic microbial ecology (CRC press, 1993).

137. A. J. Klee, A computer program for the determination of most probable number and its confidence limits. J. Microbiol. Methods 18, 91–98 (1993).

138. M. Martin, Cutadapt removes adapter sequences from high-throughput sequencing reads. EMBnet. J. 17, 10–12 (2011).

139. A. Dobin, et al., STAR: ultrafast universal RNA-seq aligner. Bioinformatics 29, 15–21 (2013).

140. S. Anders, P. T. Pyl, W. Huber, HTSeq—a Python framework to work with high-throughput sequencing data. bioinformatics 31, 166–169 (2015).

141. M. I. Love, W. Huber, S. Anders, Moderated estimation of fold change and dispersion for RNA-seq data with DESeq2. Genome Biol. 15, 1–21 (2014).

142. J. Köster, S. Rahmann, Snakemake—a scalable bioinformatics workflow engine. Bioinformatics 28, 2520–2522 (2012).

143. Z. Bar-Joseph, D. K. Gifford, T. S. Jaakkola, Fast optimal leaf ordering for hierarchical clustering. Bioinformatics 17, S22--S29 (2001).

144. J. H. Ward Jr, Hierarchical grouping to optimize an objective function. J. Am. Stat. Assoc. 58, 236–244 (1963).

145. S. Götz, et al., High-throughput functional annotation and data mining with the Blast2GO suite. Nucleic Acids Res. 36, 3420–3435 (2008).

146. C. Camacho, et al., BLAST+: architecture and applications. BMC Bioinformatics 10, 1–9 (2009).

147. S. F. Altschul, W. Gish, W. Miller, E. W. Myers, D. J. Lipman, Basic local alignment search tool. J. Mol. Biol. 215, 403–410 (1990).

148. T. Aramaki, et al., KofamKOALA: KEGG Ortholog assignment based on profile HMM and adaptive score threshold. Bioinformatics 36, 2251–2252 (2020).

149. P. Törönen, L. Holm, PANNZER—a practical tool for protein function prediction. Protein Sci. 31, 118–128 (2022).

150. A. Scheffel, N. Poulsen, S. Shian, N. Kröger, Nanopatterned protein microrings from a diatom that direct silica morphogenesis. Proc. Natl. Acad. Sci. 108, 3175–3180 (2011).

151. M. A. Maniscalco, M. A. Brzezinski, J. W. Krause, K. Thamatrakoln, Decoupling silicon metabolism from carbon and nitrogen assimilation poises diatoms to exploit episodic nutrient pulses in a coastal upwelling system. Front. Mar. Sci. 10, 1291294 (2023).

152. A. Kotzsch, et al., Biochemical composition and assembly of biosilica-associated insoluble organic matrices from the diatom Thalassiosira pseudonana. J. Biol. Chem. 291, 4982–4997 (2016).

153. E. G. Bligh, W. J. Dyer, A rapid method of total lipid extraction and purification. Can. J. Biochem. Physiol. 37, 911–917 (1959).

154. K. J. Popendorf, H. F. Fredricks, B. A. S. Van Mooy, Molecular ion-independent quantification of polar glycerolipid classes in marine plankton using triple quadrupole MS. Lipids 48, 185–195 (2013).

155. J. Hummel, et al., Ultra performance liquid chromatography and high resolution mass spectrometry for the analysis of plant lipids. Front. Plant Sci. 2, 54 (2011).

156. H. C. Holm, et al., Global ocean lipidomes show a universal relationship between temperature and lipid unsaturation. Science (80-.). 376, 1487–1491 (2022).

157. J. R. Collins, B. R. Edwards, H. F. Fredricks, B. A. S. Van Mooy, LOBSTAHS: an adduct-based lipidomics strategy for discovery and identification of oxidative stress biomarkers. Anal. Chem. 88, 7154–7162 (2016).

158. C. A. Smith, E. J. Want, G. O’Maille, R. Abagyan, G. Siuzdak, XCMS: processing mass spectrometry data for metabolite profiling using nonlinear peak alignment, matching, and identification. Anal. Chem. 78, 779–787 (2006).

159. C. Kuhl, R. Tautenhahn, C. Bottcher, T. R. Larson, S. Neumann, CAMERA: an integrated strategy for compound spectra extraction and annotation of liquid chromatography/mass spectrometry data sets. Anal. Chem. 84, 283–289 (2012).

160. S. M. Bent, et al., Lipid biochemical diversity and dynamics reveal phytoplankton nutrient-stress responses and carbon export mechanisms in mesoscale eddies in the North Pacific Subtropical Gyre. Front. Mar. Sci. 11, 1427524 (2024).

161. K. R. Heal, et al., Marine community metabolomes carry fingerprints of phytoplankton community composition. Msystems 6, 10–1128 (2021).

162. H. Wickham, H. Wickham, Getting Started with ggplot2. ggplot2 Elegant Graph. data Anal. 11–31 (2016).

163. R. Kolde, pheatmap: Pretty Heatmaps. (2025). Available at: https://cran.r-project.org/package=pheatmap.

164. T. L. Pedersen, patchwork: The Composer of Plots. (2025). Available at: https://patchwork.data-imaginist.com/.

