## Supplementary Figures for "RNA virus-mediated metabolic reprogramming in a bloom-forming, marine diatom"

**This PDF file includes:**

Figures S1 - S12

List of Data Tables S1 – S8

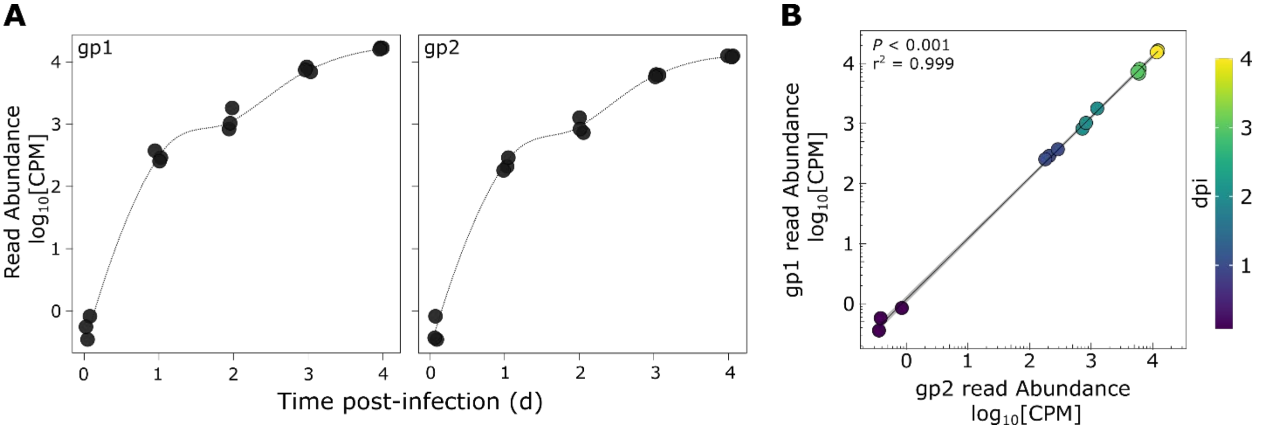


**Figure S1. CtenRNAV virus reads in RNA-seq data. A)** Mapped read abundance ($\log_{10}$[CPM]) of viral replicase polyprotein D1P48_gp1 (left) and viral structural polyprotein D1P48_gp2 (right) throughout the time course of infection in cultures infected with CtenRNAV. Biological triplicates are plotted with LOESS regression. **B)** Correlation between mapped reads ($\log_{10}$[CPM]) of CtenRNAV replicase gene gp1 (D1P48_gp1) and CtenRNAV structural polyprotein gene gp2 (D1P48_gp2) in *C. tenuissimus* throughout the time course of infection ($n = 3$). Color scale denotes days post infection (dpi). Line denotes linear regression of the data with 95% confidence interval in grey.

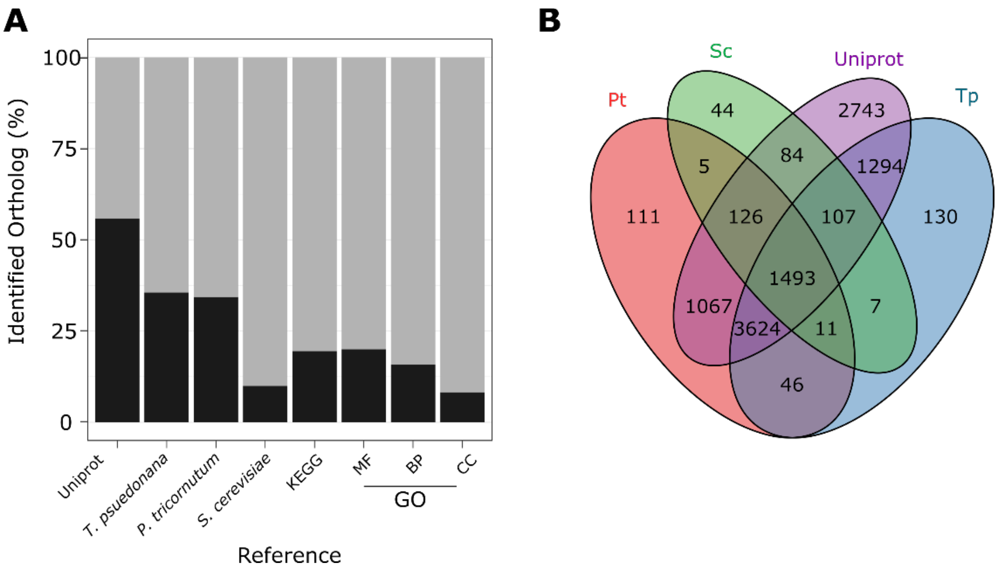


**Figure S2. Ortholog identification and function annotation summary of the *C. tenuissimus* genome.** A) Percent of predicted proteins in the *C. tenuissimus* genome with identified orthologs (black) in the Uniprot database, reference genomes (*Thalassiosira psueodonana, Phaeodactylum tricornutum* and *Saccharomyces cerevisiae*) and with KEGG assignments and Gene Ontology (GO) assignments, according to Molecular Function (MF), Biological Processes (BP) and Cellular Component (CC). B) Venn diagram depicting shared and unique orthologs in *C. tenuissimus* with *T. pseudonana* (Tp)*, P. tricornutum* (Pt) and *S. cerevisiae* (Sc) and Uniprot databases.

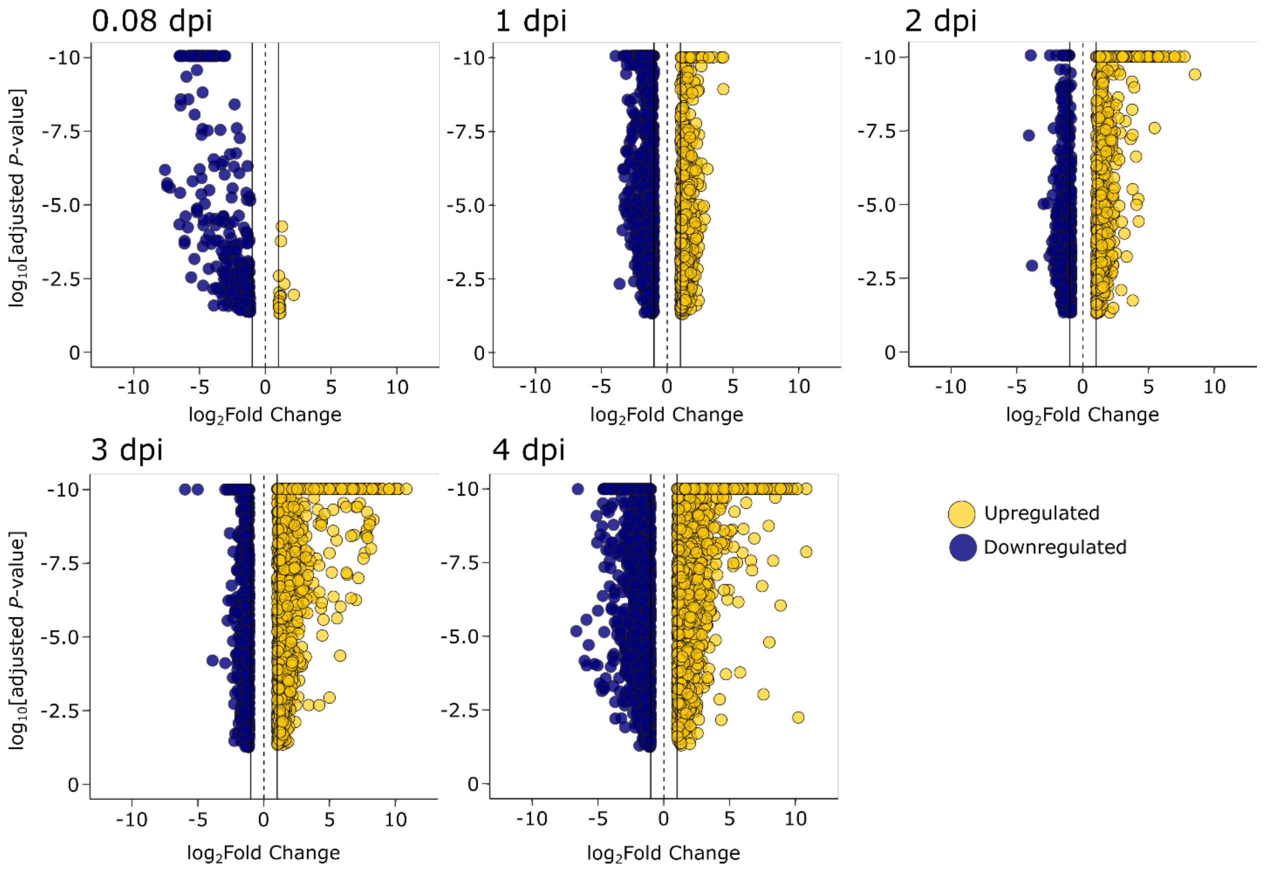


**Figure S3. Host differential expression analysis during CtenRNAV virus infection.** Volcano plots depicting differentially expressed genes in *C. tenuissimus* throughout the time course of CtenRNAV virus infection. Y axis denotes $\log_{10}$ transformed adjusted *P*-values as a function of $\log_2$ Fold Change for each gene between infected and Ctrl cultures. Yellow symbols represent genes that were upregulated in infected cells and blue symbols represent genes that downregulated in infected cells as compared to uninfected Ctrl cultures. For visualization, genes with adjusted P-values < $10^{-10}$ were placed at a *P*-value of $10^{-10}$.

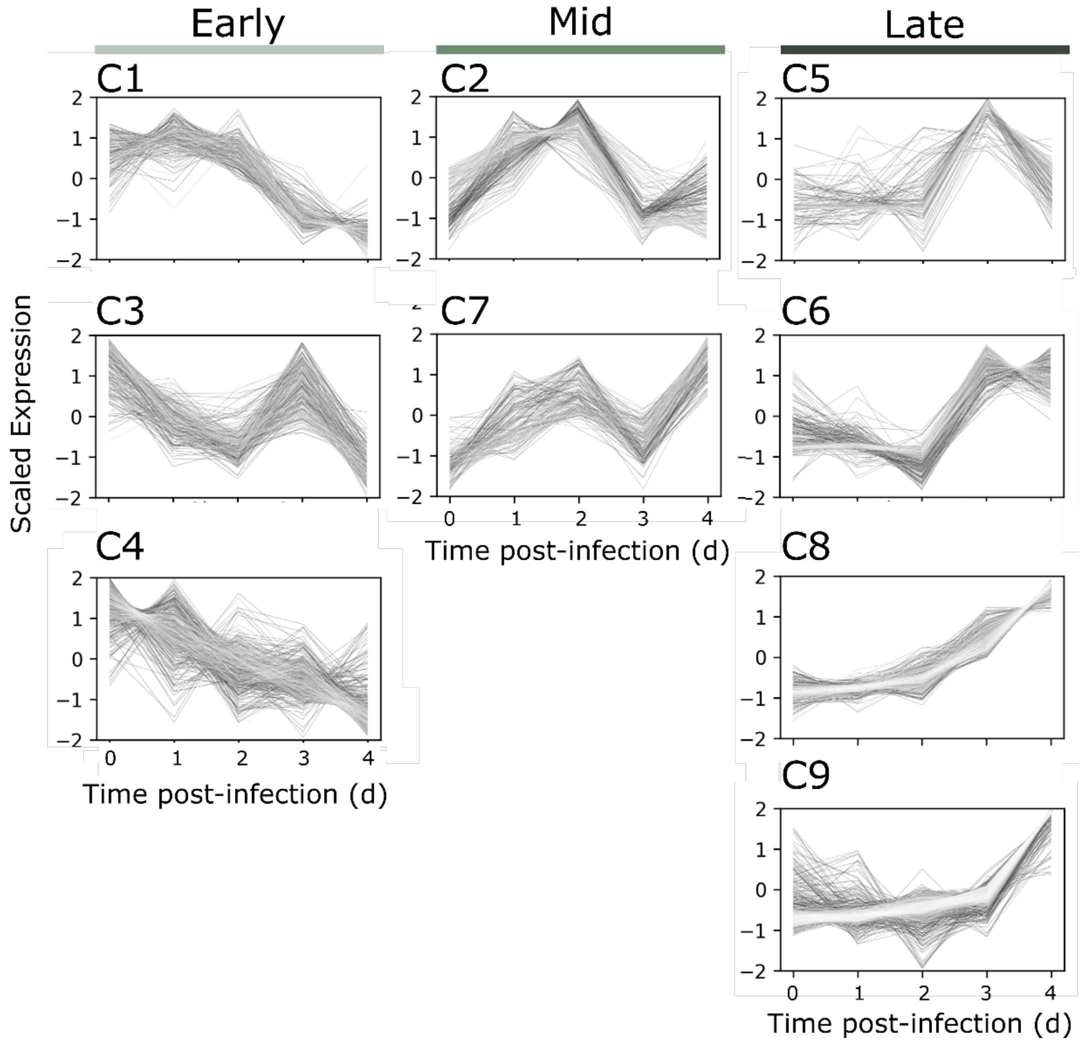


**Figure S4. Scaled expression of differentially expressed gene clusters.** Scaled expression patterns of differentially expressed gene clusters in CtenRNAV infected *C. tenuissimus* cultures throughout the time course of infection as identified by K-means analysis and organized according to expression pattern during 'Early' (light green), 'Mid' (green) and 'Late' (dark green) stage of infection.

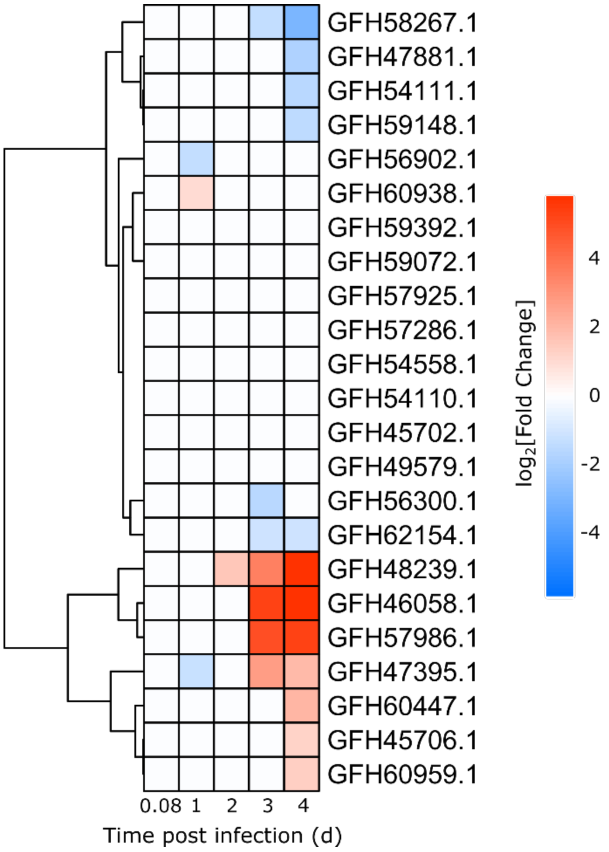


**Figure S5. Differential expression of amino acid transporters during CtenRNAV infection in *C. tenuissimus*.** Heatmap of differentially expression analysis of putative amino acid transporters in *C. tenuissimus* in CtenRNAV-infected cultures as compared to uninfected Ctrl cultures throughout the time course of infection (individual boxes). Symbol color depicts Fold Change ($log_2FC$) for each pairwise comparison. White boxes represent timepoints in which genes were not significantly differentially expressed. Full description of gene annotations, fold change and adjusted *P*-values are available in Data S2 and S3.

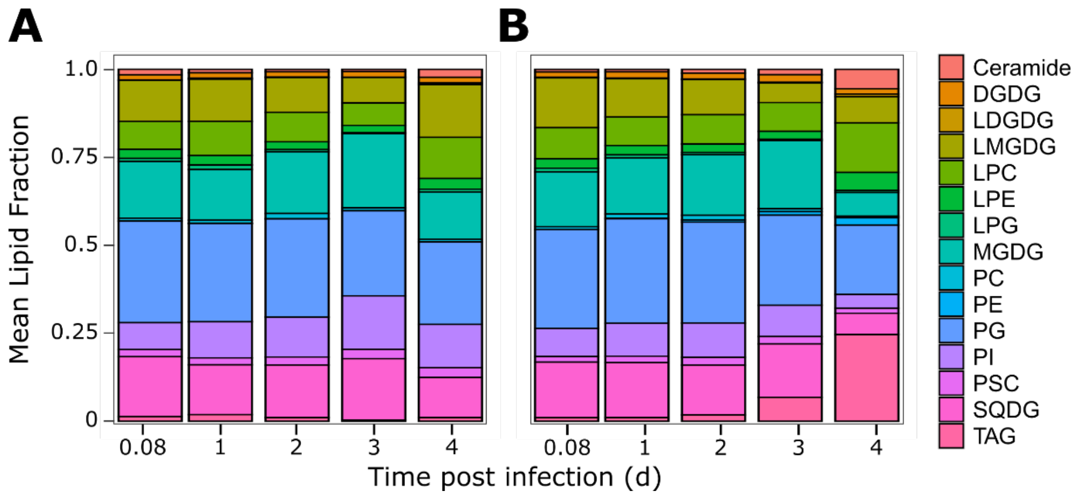


**Figure S6. Relative lipid abundance in *C. tenuissimus* throughout CtenRNAV infection.** Mean average relative abundance of intact polar lipids and triacylglycerol in **A**) uninfected Ctrl and **B**) CtenRNAV-infected cultures (n=3, except CtenRNAV-infected treatment at 0.08 dpi where 1 replicate was lost). Abbreviations are as follows: digalactosyldiacylglycerol (DGDG); monogalactosyldiacylglycerol (MGDG); phosphatidylcholine (PC); phosphatidylethanolamine (PE); phosphatidylglycerol (PG); phosphatidylinositol (PI); phosphatidylsulfocholine (PSC); sulfoquinovosyldiacylglycerol (SQDG); triacylglycerol (TAG). Lyso-polar lipid species (i.e. glycerolipids with a single acyl chain) include lyso-digalactosyldiacylglycerol (LDGDG); lyso-monogalactosyldiacylglycerol (LMGDG), lyso-phosphatidylcholine (LPC); lyso-phosphatidylethanolamine (LPE). See Data S7 for full list of annotated lipids.

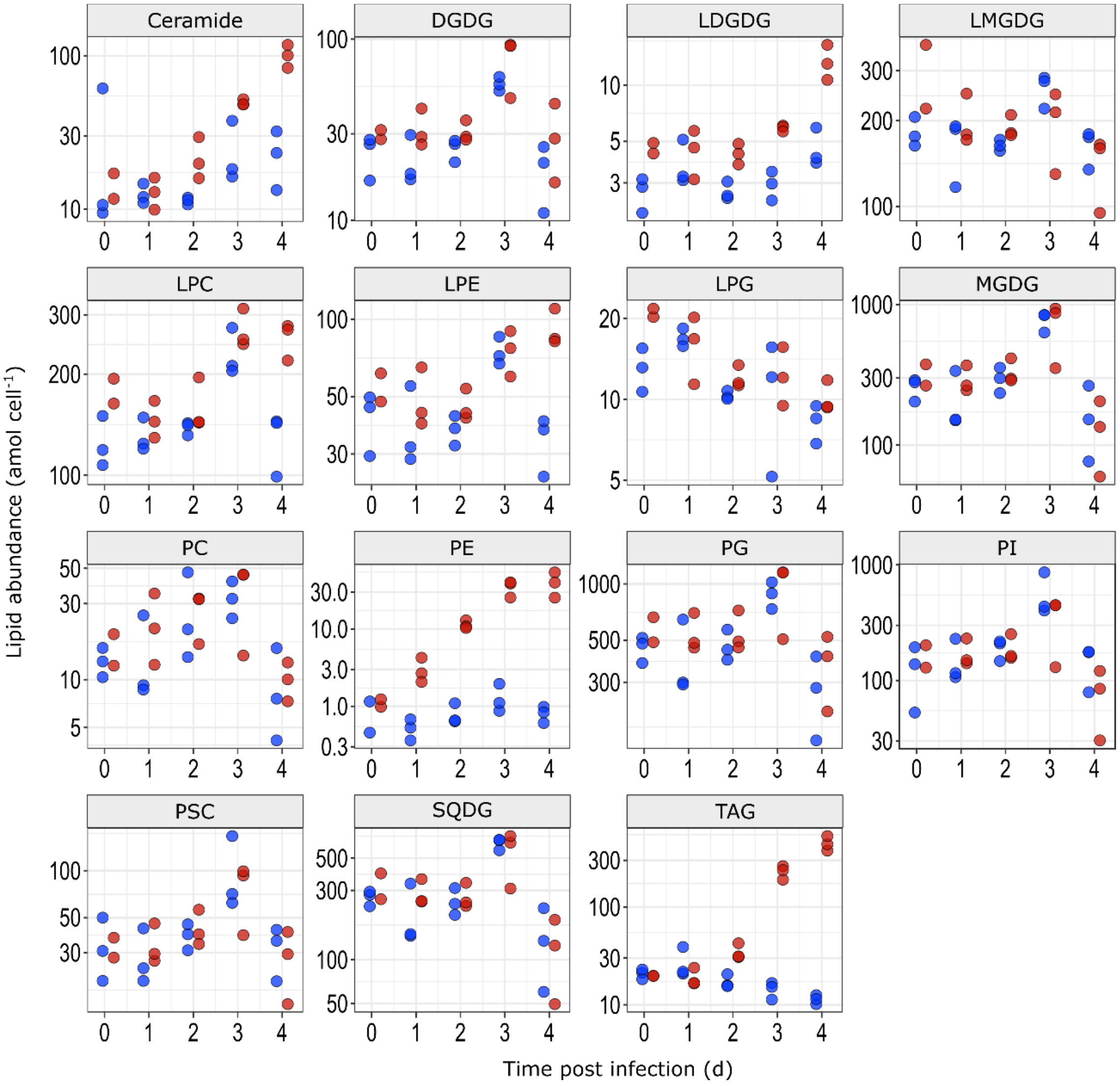


**Figure S7. Polar membrane lipids and triacylglycerol abundance in *C. tenuissimus* throughout CtenRNAV virus infection.** Cellular abundance (attomoles cell$^{-1}$) of polar membrane lipids and triacylglycerol throughout the time course of infection in CtenRNAV-infected (red) and Ctrl (blue) cultures (n= 3). Abbreviations are as follows: digalactosyldiacylglycerol (DGDG); monogalactosyldiacylglycerol (MGDG); phosphatidylcholine (PC); phosphatidylethanolamine (PE); phosphatidylglycerol (PG); phosphatidylinositol (PI); phosphatidylsulfocholine (PSC); sulfoquinovosyldiacylglycerol (SQDG); triacylglycerol (TAG). Lyso-polar lipid species (i.e. glycerolipids with a single acyl chain) include lyso-digalactosyldiacylglycerol (LDGDG); lyso-monogalactosyldiacylglycerol (LMGDG), lyso-phosphatidylcholine (LPC); lyso-phosphatidylethanolamine (LPE).

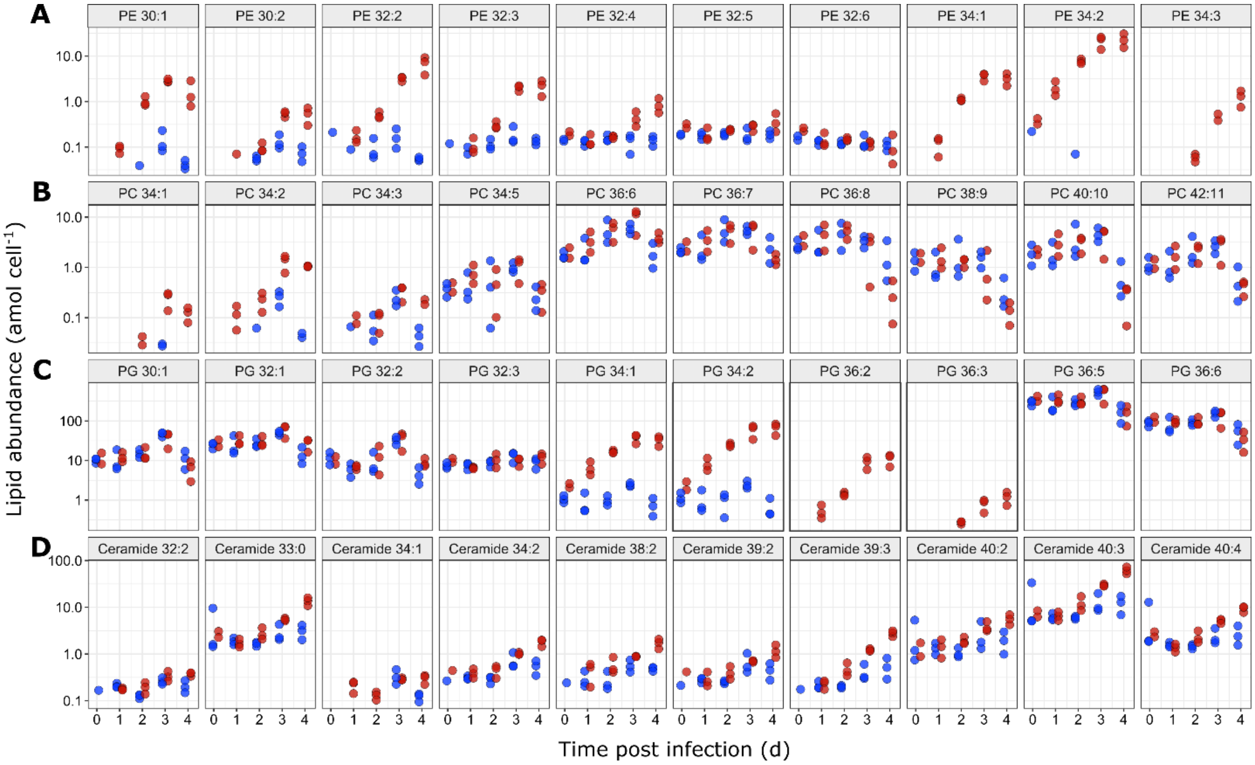


**Figure S8. Cellular abundance of major lipid class compounds in *C. tenuissimus* during CtenRNAV virus infection.** Time course of cellular abundance (attomoles $cell^{-1}$) of selected major **A)** phosphatidylethanolamine (PE) **B)** phosphatidylcholine (PC), **C)** phosphatidylglycerol (PG), and **D)** ceramide compounds in CtenRNAV-infected (red) and Ctrl (blue) cultures (n=3). Numbers in panel labels (e.g. 30:1) indicate combined acyl chain carbon:number of double bonds in a compound (or combined acyl chain and long-chain sphingosine base carbons, in the case of ceramide lipids).

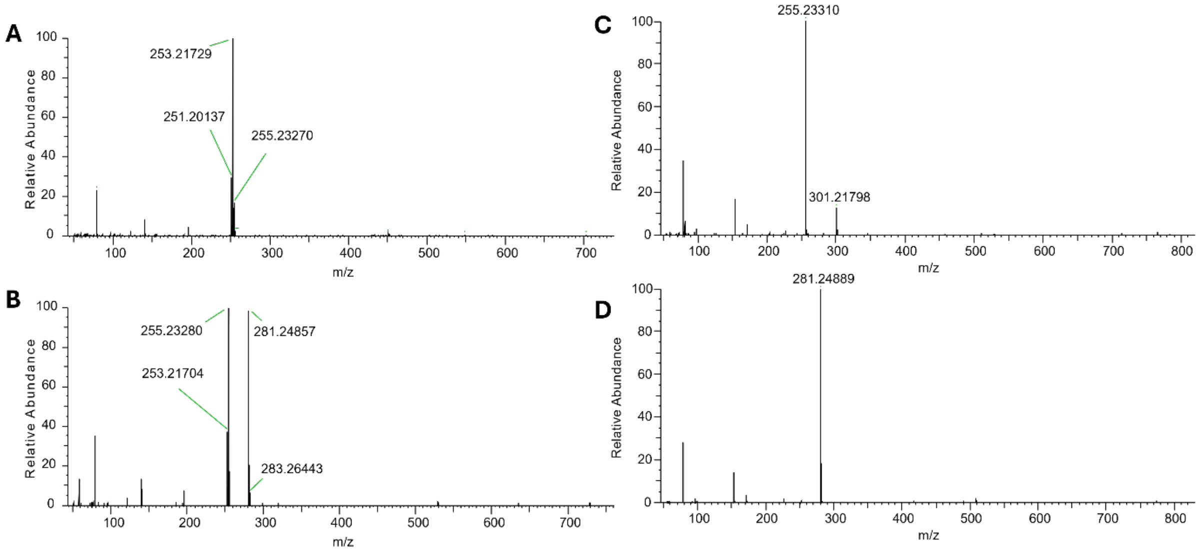


**Figure S9. Negative mode mass spectral fragmentation of typical lipid species in Ctrl and CtenRNAV-infected *C. tenuissimus*.** Negative mode mass spectral fragmentation of typical phosphatidylethanolamine (PE; panels **A** and **B**) and phosphatidylglycerol (PG; panels **C** and **D**) species in Ctrl (**A,C**) and CtenRNAV-infected (**B, D**) *C. tenuissimus*. **A)** Fragmentation of PE 32:2 (i.e. 32 acyl C, 2 double bonds), with fragments m/z 251, 253, 255 indicating free fatty acids (FFA) 16:2, 16:1, and 16:0, implying PE 32:2 is comprised of PE 16:0/16:2 and PE 16:1/16:1. **B)** Fragmentation of PE 34:1 in CtenRNAV-infected cells, with primary fragments 255 and 281 indicating PE 34:1 is primarily composed of FFA16:0 and 18:1, with minor contributions of FFA 16:1 and 18:0, fragments 253 and 283. **C)** Fragmentation of PG 36:5 (i.e. 36 acyl C, 5 double bonds, the most abundant PG compound in Ctrl *C. tenuissimus*), with fragments m/z 255 and 301 indicating PE 36:5 is composed nearly entirely of the saturated FFA 16:0 and polyunsaturated FFA 20:5. **D)** Fragmentation of PG 36:2, with primary fragment m/z 281 indicating PG 36:2 is primarily composed of FFA 18:1.

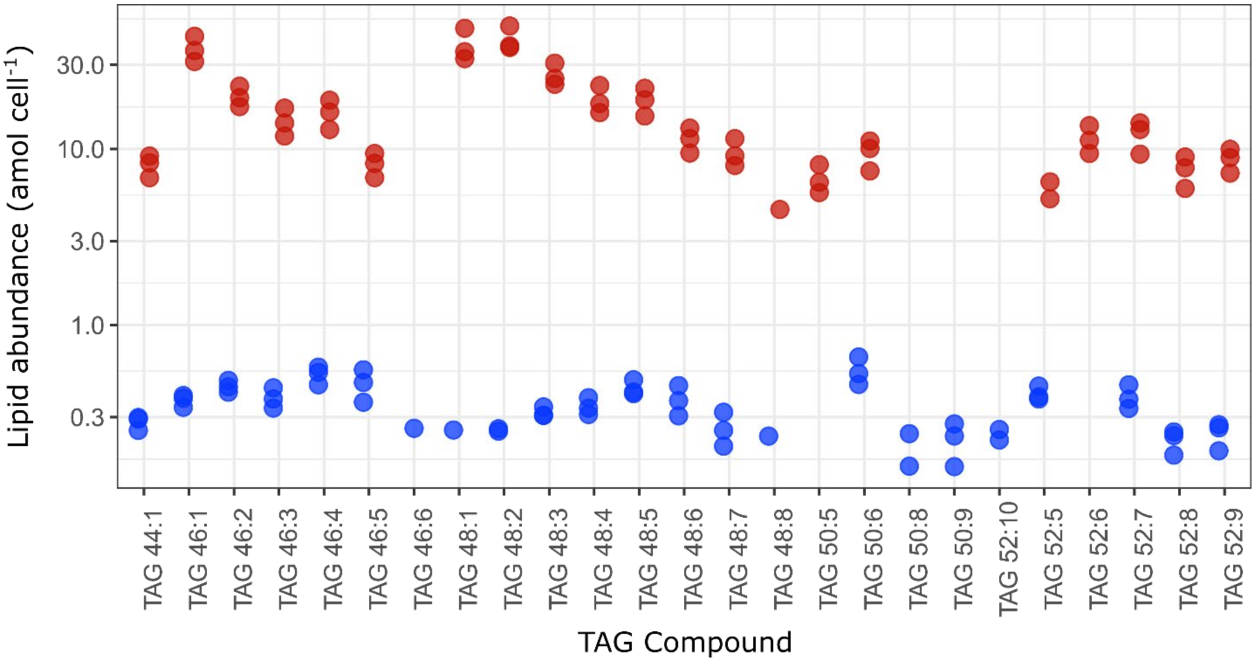


**Figure S10**. **Triacylglycerol compound abundances during CtenRNAV**. Abundances of individual triacylglycerol (TAG) lipid molecules (attomoles cell$^{-1}$) in CtenRNAV-infected (red) and uninfected, Ctrl (blue) cultures (n = 3 biological replicates). Number labels, e.g. TAG 48:10, indicate 48 carbon atoms and 10 double bonds shared among three acyl chains.

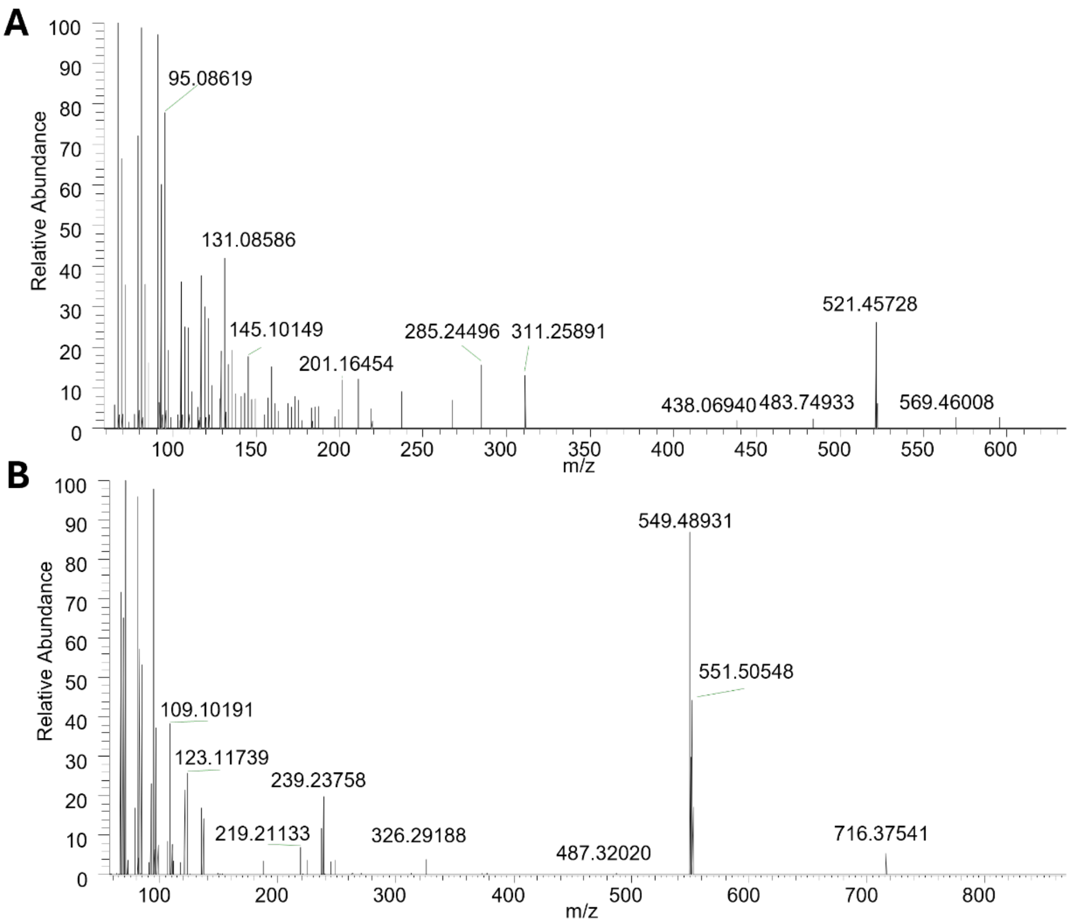


**Figure S11**. Positive mode mass spectral fragmentation analyses of prominent triacylglycerol (TAG) molecules in **A)** Ctrl and **B)** CtenRNAV-infected cultures. **A)** An MS2 spectrum of TAG 50:6 (indicating 50 carbons and six double bonds between three acyl chains; m/z 840.7075669) in an uninfected Ctrl cells, with fragments m/z 285, m/z 311, and m/z 521 (neutral loss of 319 Da) indicating fatty acids 16:0, 18:1, and 20:5; and **B)** an MS2 spectrum of TAG 48:1 (m/z 822.7545172) in a CtenRNAV-infected culture, with fragments m/z 549 and m/z 551(neutral losses of 273 and 271 Da), indicating loss of fatty acids 16:0 and 16:1.

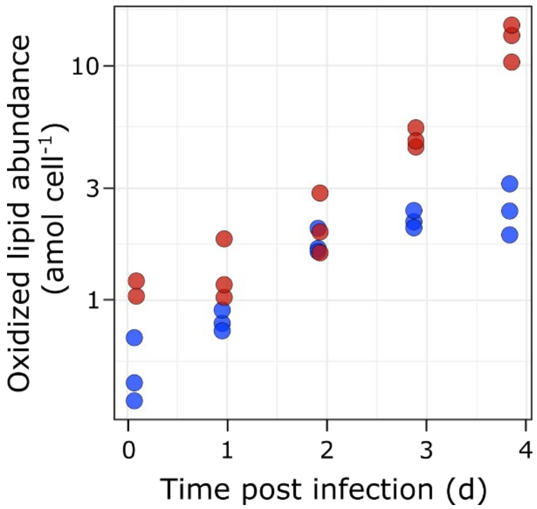


**Figure S12. Oxidized and hydroperoxidized lipids during CtenRNAV infection in *C. tenuissimus.*** Total quantified oxidized and hydroperoxidized lipids (attomoles cell$^{-1}$), including membrane lipids and triacylglycerol in CtenRNAV-infected (red) and uninfected, Ctrl (blue) cultures throughout the time course infection (n = 3 biological replicates).

### Supplementary Data Files

**Data S1**. RNA sequencing summary

**Data S2.** Ortholog identification and functional annotation in *C. tenuissimus*

**Data S3.** Differentially expressed genes in *C. tenuissimus* during CtenRNAV infection.

**Data S4.** Scaled expression and cluster analysis of differentially expressed genes in CtenRNAV-infected *C. tenuissimus*

**Data S5.** Gene Ontology enrichment analysis

**Data S6.** Annotation of silicon metabolism related genes in *C. tenuissimus*

**Data S7.** Particulate lipid compounds identified throughout CtenRNAV infection in *C. tenuissimus*

**Data S8.** Relative and cellular abundance of lipids groups identified throughout CtenRNAV infection in *C. tenuissimus*
